# Comprehensive and accurate genome analysis at scale using DRAGEN accelerated algorithms

**DOI:** 10.1101/2024.01.02.573821

**Authors:** Sairam Behera, Severine Catreux, Massimiliano Rossi, Sean Truong, Zhuoyi Huang, Michael Ruehle, Arun Visvanath, Gavin Parnaby, Cooper Roddey, Vitor Onuchic, Daniel L Cameron, Adam English, Shyamal Mehtalia, James Han, Rami Mehio, Fritz J Sedlazeck

## Abstract

Research and medical genomics require comprehensive and scalable solutions to drive the discovery of novel disease targets, evolutionary drivers, and genetic markers with clinical significance. This necessitates a framework to identify all types of variants independent of their size (e.g., SNV/SV) or location (e.g., repeats). Here we present DRAGEN that utilizes novel methods based on multigenomes, hardware acceleration, and machine learning based variant detection to provide novel insights into individual genomes with ∼30min computation time (from raw reads to variant detection). DRAGEN outperforms all other state-of-the-art methods in speed and accuracy across all variant types (SNV, indel, STR, SV, CNV) and further incorporates specialized methods to obtain key insights in medically relevant genes (e.g., HLA, SMN, GBA). We showcase DRAGEN across 3,202 genomes and demonstrate its scalability, accuracy, and innovations to further advance the integration of comprehensive genomics for research and medical applications.

## Introduction

Over the last decade, the advent of genomic sequencing as a common methodology in genomics, genetics, and medical applications has enabled multiple discoveries and insights for diseases, population diversity, evolutionary mechanisms, and personalized medicine strategies^1–4^. This was in large part possible due to improvements in next-generation sequencing (NGS) (i.e., Illumina) in terms of costs, high data quality, and scalability^1^. Highly accurate methods for the detection of single nucleotide variations (SNV) and smaller (<50bp) insertions or deletions (indel) have been at the forefront of variant detection and interpretation. Despite the amount of attention SNV have garnered, they are not the only variant type that differentiates two genomes^5,6^. Recently, an increasing number of studies incorporate structural variation (SV)^7–9^ into their analysis. SVs are often defined to be 50bp or larger and lead to deletions, insertions, amplifications, or rearrangements of a genome^7^. Copy number variation (CNV) is another genomic variation that arises from deletions (loss of copies) or duplications (gain of copies) of a specific DNA segment^7^. Another understudied variant type are tandem repeat expansions that are mainly defined by their low sequence entropy/complexity^10,11^. These types of variants have been associated with many diseases, diversity, and evolutionary patterns. The detection and interpretation of them remain challenging, but multiple specialized methods have been proposed^5,7^. While all these variant types are present across genomes, many studies often focus on only SNV or subsets of variant types independently due to the challenges of joint detection and accurate reporting of these variant classes. Additional challenges arise from highly diverse and repetitive regions of the genome that further complicate the analysis^6,12^. While these variant types likely interact together, these relations are lost when analyzed independently. Thus, more comprehensive approaches that can scale are required.

One proposed way to unify variant discovery is via specialized sequencing technologies, i.e., long reads, that have been reported to improve certain aspects such as SV detection (e.g., Oxford Nanopore Technologies (ONT) or PacBio)^5,7^. These technologies have matured significantly over the past few years and are becoming more commonly available^5^. However, long read technologies are still often limited by their costs, data quality, and scalability and more often by their sample requirements in terms of quantity and quality^5^. This often hinders their application across larger populations or even legacy samples. Interestingly what these sequencing technologies have demonstrated is that the alleles that are identifiable using their long reads are indeed also often present and identifiable in short reads^13^. This has been most successfully shown in SV genotyping using graph genomes^13,14^. Recent improvements including graph genome approaches have been shown to improve SV genotyping and the mapping of short reads^15^. Nonetheless, these methods often pose challenges to apply them at scale or generate comprehensively and thus have often been applied to only re-identify certain alleles (i.e., genotyping)^16^ making their utility so far very limited^15^. Single improvements need to act together to fully detangle the complex genomic landscape of an individual, even more so on a population scale.

The current trend is often not only to identify and interpret variants in only the coding regions of the genome, but as well investigate the impact of variants across the entire genome using whole genome sequencing, which further adds to the complexity of the challenges due to repetitiveness (e.g., segmental duplications) complex polymorphisms and annotations^6,17^. The central question to addressing these challenges is what is needed to improve the interpretation of all variant types in order to identify novel candidate disease alleles or genes. To tackle this, the typical approach is to increase the number of samples that are analyzed to obtain robust population allele estimations. This motivates multiple large scale studies (e.g. Centers for Common Disease Genomics (CCDG)^18^, Trans-Omics for Precision Medicine (TOPMed)^19^, All of US (AoU), UK Biobank (UKBB)) focusing on Illumina sequencing which substantiates short reads’ role as the workhorse of genomics and genetics. It also requires a scalable and unified software framework to comprehensively identify all variant types (SNV, indel, SV, and repeat expansion), which has not been realized so far^6^. A framework capable of this would not only scale the identification of the variation landscape from a single genome to millions of genomes, but would enable us to obtain novel and key insights into multiple adult diseases that are currently poorly understood due to a sole focus on SNV^20,21^.

Here we present new developments of Dynamic Read Analysis for GENomics (DRAGEN) and its optimization in SNV and indel calling as well as its ability to detect the entire landscape of variations (CNV, SV, repeat expansions, specialized methodologies for certain regions: HLA, SMN1&2, etc.). These developments bring together advancements in genomic algorithm development to address longstanding issues of scalability, accuracy, and comprehensiveness of variant detection across all sizes and types of alleles and thus fully resolve individual genomes at scale and cost. The accuracy of DRAGEN is boosted by the first multigenome (graph) implementation that scales and enables the detection of variant types beyond just SNV. In this study, we introduce and benchmark DRAGEN’s 14 sub components (SNV, SV, STR, CNV, nine targeted callers including four new callers, and gVCF genotyper) and illustrate their ability to scale across large cohorts by analyzing the 1000 genome project (1kGP)^22^. We reveal new insights into the diversity of genome across population with a special focus on medically relevant genes to demonstrate the genomic and medical utility of DRAGEN. We introduce novel methods to compare and merge the variants produced to further emphasize DRAGEN’s ability to analyze multiple variant classes together. This includes novel SNV and indel merging strategy to scale and produce fully genotyped population variant call format (VCF) files. Similarly, we provide novel solutions to combine STR, SV, and CNV into one population VCF file. Both methods allow, for the first time, the handling of all variant types together and promote the assessment of large variants for cohort studies. We demonstrate this across 3,202 whole genome samples from the 1kGP cohort. This work demonstrates DRAGEN’s capability to solve the current issues and limitations of research and clinical genomics to further the discovery of novel disease targets ranging from common to rare disease studies and novel insights into the diversity of genomes all together.

## Results

### Novel algorithms to enable comprehensive genomics at scale and accuracy

In this manuscript, we present a novel framework (DRAGEN version 4.2.4) to identify all types of genomic variations at scale and cost. **Figure 1** gives a brief overview of DRAGEN’s main components. First, each sample is mapped to a multigenome (graph) consisting of a reference and several assemblies e.g., GRCh38 in addition to 64 haplotypes (32 samples) together with reference corrections previously reported^24^ to overcome errors on the human genome. The multigenome (graph) includes variants from multiple genome assemblies to better represent sequence diversity between individuals throughout the human population. In brief, the seed based mapping considers both primary (e.g., GRCh38) and secondary contigs (phased haplotypes from other populations) throughout the multigenome. The alignment is controlled over established relationships of primary and secondary contigs and is adjusted accordingly for mapping quality and scoring (see **methods** for details). DRAGEN’s mapping process for a 35x whole genome sequence (WGS) paired end data set, requires approximately 8 minutes of computation time using an onsite DRAGEN server (**Supplementary Table S1** has details of time taken in each step for both AWS F1 instance and onsite Phase4 server). The multigenome can be updated with advancements (e.g., T2T-CHM13 or HPRC pan-genome reference) and can enable a more precise and comprehensive alignment of the short reads. These improved alignments are leveraged for variant calling.

**Figure 1:**
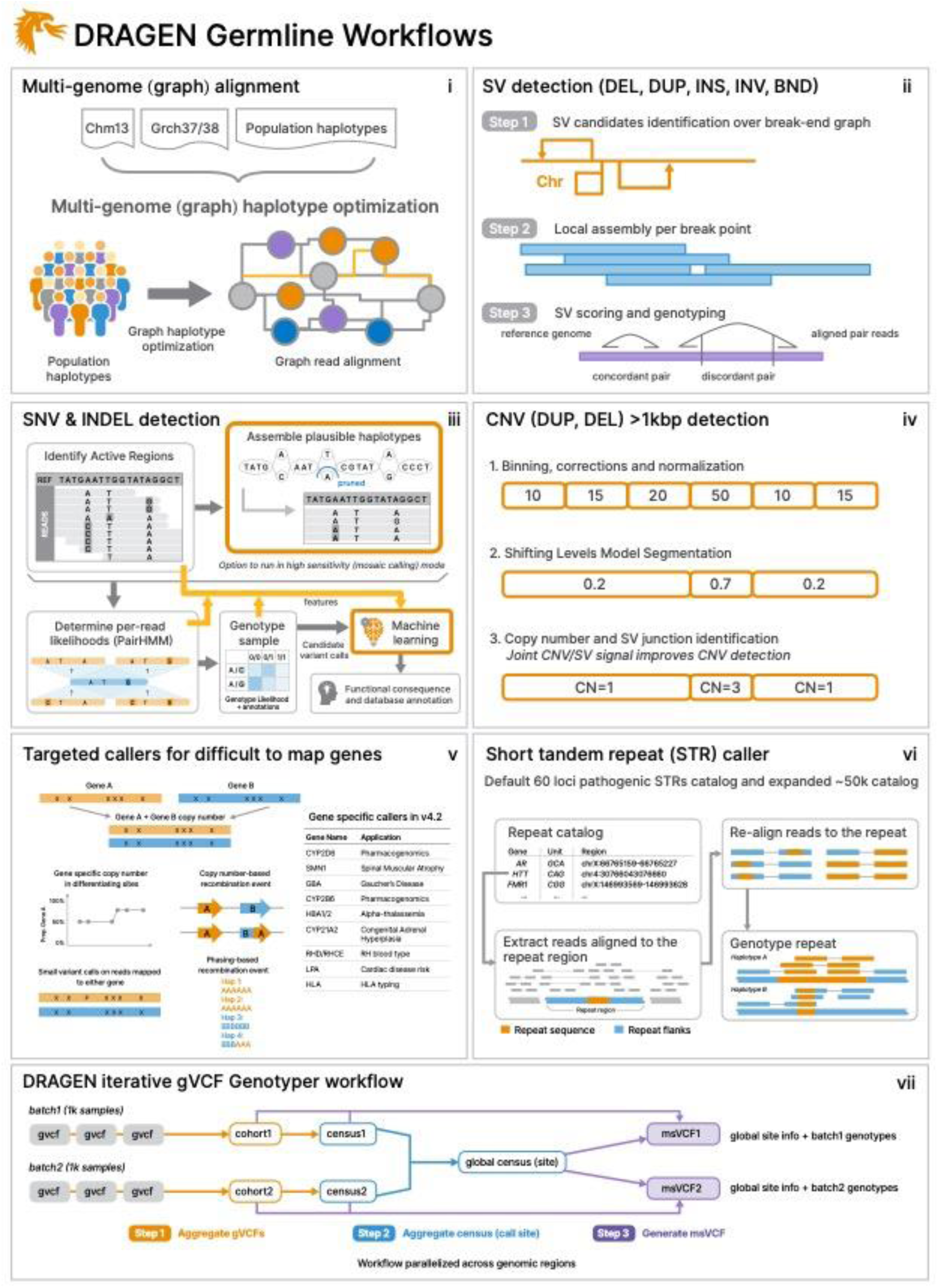
Overview of DRAGEN variant calling pipeline. DRAGEN improves the variant identification from single bp up to multiple Mbp of alleles. This is achieved by implementing multiple optimized novel concepts: i) Mapping utilizes a multigenome (graph) including 64 haplotypes; ii) SV calling is significantly improved over local assemblies based on breakpoint graphs; iii) SNV calling is improved using multiple novelties including machine learning based scoring and filtering; iv) CNV calling utilizes the multigenome (graph) and the SV calling information to make informed decisions; vi) Additional nine tools targeting specific difficult regions of the genome are included, four of them not reported before; vi) STR calling is integrated based on Expansion Hunter^23^; and vii) A gVCF genotyper implementation to provide a population level fully genotyped VCF file.

To identify single nucleotide variants and indels (<50bp), DRAGEN pre-assembles regions of variants using a *de Bruijn* graph, which is then input to a Hidden Markov Model (HMM) with previously estimated noise and error levels per sample. The output is a (g)VCF file. The SNV caller has key innovations to deal with noise or sequencing errors including i) sample-specific Polymerase Chain Reaction (PCR) noise estimation; ii) correlated pileup errors estimation; iii) consideration of overlapping candidate events; and iv) local assembly failures and incomplete haplotype candidates. After the initial variant calling, a machine learning (ML) framework rescores calls to further reduce false positive small variants (both SNV and indel) and recover wrongly discarded false negatives (see Figure 1 and **methods** for details).

Simultaneously, DRAGEN identifies Structural Variations (SV) (>=50bp genomic alterations) as well as copy number events (>=1kbp genomic alterations) using two methods (see Figure 1 and **methods** for details). For SV calling, DRAGEN extends Manta^25^ by introducing key concepts that significantly improve SV calling: i) new mobile element insertion detector for large insertion calling; ii) optimization of proper pair parameters for large deletion calling; iii) improved assembled contig alignment for large insertion discovery; iv) refinements to the assembly step; v) refinements in read likelihood calculations step; vi) improved handling of overlapping mates; vii) improved handling of clipped bases; and viii) filtering and precision improvements (see Figure 1 and **methods** for details). For CNV calling, DRAGEN targets 1kbp and larger variants that cause an amplification or deletion of genomic segments. This CNV caller utilizes a modified Shifting Levels Model, which identifies the most likely state of input intervals through the Viterbi algorithm (see Figure 1 and **methods** for details). The CNV caller was also designed to take into consideration the discordant and split read signals from the SV calling to be able to detect events down to 1kbp. Furthermore, DRAGEN identifies short tandem repeat mutations and analyzes known pathogenic genomic regions using a method primarily based on ExpansionHunter^23^.

Some important genes are challenging to genotype due to their high sequence similarity to pseudogenes, repetitive regions, and polymorphic nature. To overcome these challenges, DRAGEN integrates nine targeted callers for accurate genotyping of clinically relevant genes (*CYP2B6, CYP2D6, CYP21A2, GBA, HBA, LPA, RH, SMN, and HLA*), of which six of the callers are described here for the first time^26–29^. In general, DRAGEN utilizes common SNV in the population to distinguish gene targets from their paralogous copies to provide copy number estimation for each haplotype. Furthermore, DRAGEN identifies reads that do not follow the general phasing patterns and reports the recombination events that lead to these reads per sample (See **methods** for details on each caller). The *CYP2D6* and *CYP2B6* genes are important for pharmacogenomics and encode an enzyme that is responsible for metabolizing most of the commonly used drugs^30^. The recombination of gene and pseudogene can lead to deletions of part of each copy, generating gene-pseudogene fusions. The variants across *CYP21A2* can lead to Congenital Adrenal Hyperplasia ^31^*. GBA* is an important target gene due to variants that increase risk for Parkinson’s and Gaucher’s disease and Lewy body dementia^32,33^. The gene resides in a segmental duplication of 10kbp with a pseudogene *GBAP1*. The high sequence homology in *GBA/GBAP1* drives homologous recombination and can result in pathogenic gene conversions or copy number variants. The HLA genes encode proteins crucial for immune regulation and response, holding immense importance in research related to autoimmune diseases, organ transplantation, and cancer vaccines and immunotherapy^34,35^. DRAGEN includes a specialized caller to identify the HLA class I (HLA-A, -B and -C) and class II (HLA-DQA1, -DQB1, -DRB1) alleles. Mutations in the *HBA* genes (*HBA1* and *HBA2*) cause alpha thalassemia, an inherited blood disorder characterized by lowered levels of alpha globin, a fundamental building block of hemoglobin^36^. Recurrent homologous recombination can result in 3.8kbp deletions that create a hybrid copy of *HBA1* and *HBA2*, 4.2kbp deletions that delete regions that include the *HBA2* gene, or complete deletion of both. Small pathogenic variants also can be detected within *HBA*. The *LPA* gene includes a 5.5 kbp region (*KIV-2)* whose copy numbers (between 5 to 50+) are inversely related to the cardiovascular risk^37^. DRAGEN can report phased copy number for this region^29^. For *RHD/RHCE* (RH blood type), copy number predictions can be used to assess the risk of Rh allosensitization^38^. Another integrated caller identifies copy number variants across *SMN1*&*2* which can indicate Spinal Muscular Atrophy ^27^.

The genome wide simultaneous assessment for SNV, indel, STR, SV, and CNV together with reporting the results from these specialized callers takes ∼30 minutes of computation time with an onsite DRAGEN server for a 35x WGS sample. This results in a gVCF file for SNV and indels, a VCF file for each STR, CNV, and SV calls, and tabular formats for the specialized gene callers (Figure 1).

Thus, the DRAGEN pipeline is able to capture the entire range from single variants to larger variations across the entire genome at scale and reports them in standardized VCF files. The algorithms are described in detail in the methods section. This pipeline produces the most comprehensive collection of accurate variations across a human genome and has the ability to scale.

### Resolving the complete variant spectrum at scale and accuracy

We applied DRAGEN to the HG002 sample, for which multiple benchmarks are available^16,39–42^. We identified variants using DRAGEN across a 35x coverage HG002 Illumina NovaSeq 6000 2×151bp paired-end read data set (see methods). Figure 2A shows the distribution of all small and large variants across the HG002 sample and highlights the ability of DRAGEN to capture the entire variant spectrum. This resulted in ∼4.96M small variant calls that includes 4,003,042 single-nucleotide variants (SNVs) with a transition-to-transversion (Ti/Tv) ratio of 1.98 and an SNV heterozygous to homozygous (HET/HOM) ratio of 1.57. A total of 967,735 small insertions or deletions (indel) were discovered with an insertion to deletion) ratio of 1.00 and HET/HOM ratio of 1.855. For structural variants (SVs), 14,506 variants (>=50bp) were identified with 5,901 deletions (DEL), 7,174 insertions (INS), 42 duplications (DUP), 153 inversions (INV), and 616 translocations (TRA). Additionally, 1,156 copy number variants (CNVs) were identified ranging from 1kbp to 445kbp with a deletion-to-duplication ratio of 4.25. DRAGEN detects short tandem repeat (STR) expansions in 50,069 polymorphic loci including 60 pathogenic loci (homozygous reference 0/0: 37.33%, heterozygous 0/1: 27.36%, homozygous alternate 1/1: 17.8%, and heterozygous genotype composed of two different ALT alleles 1/2: 17.5%). Relative to GRCh38, 46.66% (14,636) of HG002 STRs have at least one more copy and 53.34% (16,734) have at least one less copy. Thus highlighting all the variant complexities a single genome represents.

**Figure 2:**
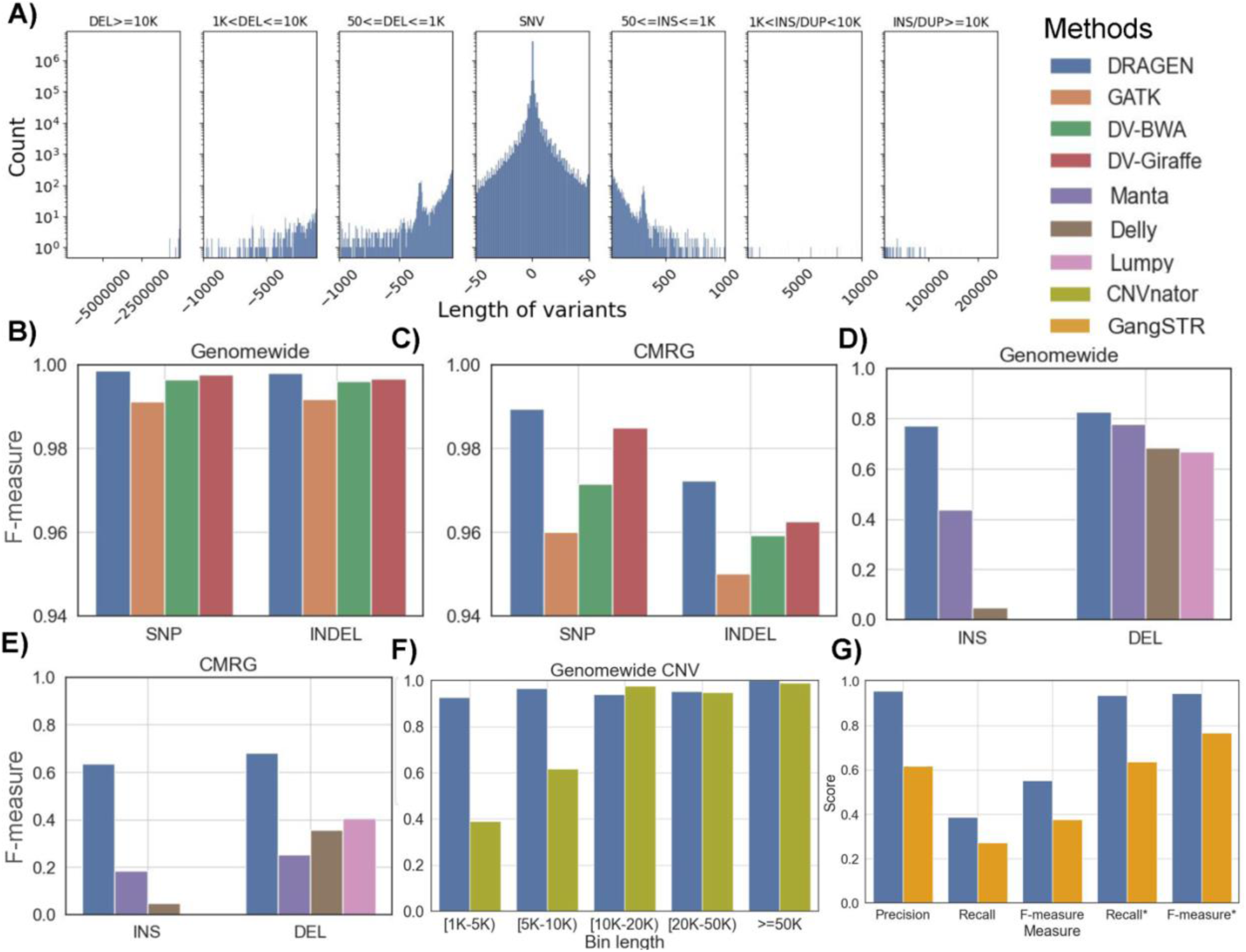
Performance overview of DRAGEN based on GIAB benchmarks. **A)** length distribution of small and large variants discovered by DRAGEN (bin sizes used for the plot (from left to right) are: 500, 250,150,50, 150, 250, 500), **B)** SNV comparison based on GIAB SNV 4.2.1, **C)** SNV call comparisons based on CMRG v1.0, **D)** Comparison of SV call performance (INS and DEL types) based on GIAB SV v0.6, **E)** Comparison of CMRG SV call performance (INS and DEL types) based on GIAB CMRG SV v1.0, **F)** CNV caller comparison of DRAGEN compared to CNVnator across different sizes of deletions based on GIAB SV v0.6, and **G)** The benchmarking of short tandem repeats using GIAB v1.0. The recall and F-measure was calculated using GIAB catalog and the recall* and F-measure* were calculated using the catalogs of DRAGEN and GangSTR.

Using these results, all the variants were evaluated against the Genome in a Bottle (GIAB) benchmarks and compared to other short read based callers (see **methods**). For SNV and indel, benchmark version 4.2.1 was used on GRCh38^40^, but for the SV benchmark (version 0.6)^39^ DRAGEN was run on a GRCh37 version of the multigenome (graph) reference. Later, the benchmark is expanded across the challenging medically relevant gene catalog ^41^ (see **methods** for details). Overall benchmarks DRAGEN demonstrates higher accuracy and impressive speed up of the analysis from raw reads to finalized variant calls within 30 min total which is better than any other existing workflow.

We first focused on SNV and indel calling for HG002 and compared its performance to other short read based methods^43^ (GATK^44^, DeepVariant^45^ with BWA^46^). We further benchmarked the recent pan genome approach: Giraffe^15^. Figure 2B shows the F-measures across SNV and indel results (**Supplementary Table S2** for details). Overall, we observed a clear advantage of DRAGEN SNV identification accuracy relative to other methods. For the overall genome-wide small variant calls, DRAGEN achieved a F-measure of 99.85% yielding a total of 11,116 errors (2,735 FPs and 8,722 FNs). Compared to DRAGEN, we observe 2.42 times more errors for DeepVariant+BWA calls (F-measure: 99.64%, 3,695 FPs, and 24,090 FNs), 1.74 times more errors for DeepVariant+Giraffe calls (F-measure: 99.74%, 4,980 FPs, and 15,021 FNs), and 5.91 times more errors for GATK+BWA calls (F-measure: 99.13%, 38,622 FPs, and 29,163 FNs) with the same Illumina sample. This is in part due to the novel methodologies implemented in the SNV calling and in the subsequent machine learning filtering (see methods). We observed improvements for substitutions and indel (2-50bp) variant types. DRAGEN achieved a higher F-measure of 99.86% (substitutions) and 99.80% (indel) compared to GATK+BWA, DeepVariant+BWA, and DeepVariant+Giraffe (**Supplementary Table S2**). Thus clearly, DRAGEN shows an improved performance on SNV and indel across the entire spectrum, improving the characterization across samples at scale.

Next, we assessed the performance of variant calling in the challenging medically relevant genes (CMRG) catalog. This GIAB benchmark spans 273 medically relevant genes that are highly repetitive and diverse and were therefore excluded from the genome wide benchmark^12^. Many of these medically relevant genes overlap segmental duplications and other challenging properties. There is interest to see if short read sequencing can effectively be used for detecting variants in these repetitive regions. Moreover, several of these medically relevant genes (e.g., *KCNE1, CBS, CRYAA, KCNJ18, MAP2K3, KMT2C,* etc.) are wrongly represented in the GRCh38 reference due to false duplication and collapsed sequence errors^24^. Corrections to these errors have been incorporated into DRAGEN variant calling. Figure 2C shows the results of the individual SNV callers with respect to F-measure (see **Supplementary Table S2** for detail evaluations). For both SNV and indel calls, DRAGEN (F-measure: 98.64%) was better than GATK (95.84%), DeepVariant+BWA (97.32%), and Deepvariant+Giraffe (98.10%). These improvements are present in both substitutions and indels (**Supplementary Table S2).** Thus outperforming the other methods with 13,931 variants genomewide and 509 variants in CMRG regions which are only identifiable by DRAGEN. We further investigated if this performance differed between exonic and intronic regions. For the exonic regions, DRAGEN achieved an F-scores of 99.78%. For intronic and intergenic regions, the achieved F-scores were 99.87% and 99.85%, respectively. Similarly, the variant calling performance was evaluated on exon and intron regions using the GIAB CMRG benchmark set. DRAGEN achieved F-scores of 98.97% and 98.66% on exon and intron regions, respectively.

In addition to the clear improvements of DRAGEN for SNV (Figure 2B-C), its performance across SV (>50bp) was also improved. The DRAGEN results were compared with SV calls reported by Manta^25^, Delly^47^, and Lumpy^48^ (Figure 2D-E) (see methods for details). For insertions, which are often the hardest for SV callers^7^, DRAGEN achieved an F-score of 76.90%, which more than doubled the performance of Manta (34.90%) and Delly (4.70%) (Lumpy didn’t report any insertions). This is due to multiple algorithmic innovations in DRAGEN (see **Methods**). Similarly, DRAGEN achieved a better F-score (82.60%) for deletion (50bp+) compared to Manta (70.80%), Delly (68.30%), and Lumpy (66.80%). **Supplementary Table S3** contains details across the SV variant types. DRAGEN performance was also compared for SV detection on the challenging medically relevant gene (CMRG) regions. DRAGEN again outperformed the other variant callers with F-measures of 63.50% and 68.00% for INS (Figure 2D) and DEL (Figure 2E) types, respectively. This showcases the ability of short reads to detect SV with high accuracy even in repetitive regions.

DRAGEN also reports copy number variations (CNV), which includes larger deletions and duplications (see methods). Here CNV are adjusted for the called SV to improve breakpoint accuracy where possible (**see Methods**). The performance was compared against CNVnator^52^ copy number discovery tool and benchmarked using the >1kbp DEL SV records from GIAB SV benchmark set (shown in Figure 2F). For CNVs with length in the range of 1-5kbp and 5-10kbp, DRAGEN performed significantly better with F-measures of 92.60% (vs 39.20% CNVnator) and 96.60% (vs 61.80% CNVnator), respectively. For CNVs with lengths in the range 10-20kbp, 20-50kbp and >50kbp, similar performance by DRAGEN (F-measures 94.10%, 95.20%, 100.00% respectively) and CNVnator (97.60%, 94.90% and 99.00% respectively) was observed. **Supplementary Table S4** contains all the benchmarking results.

Similar to SV, short tandem repeats (STR) are often challenging to resolve due to their repetitiveness and complexity^49^. The accuracy of STR detection by DRAGEN was evaluated using the GIAB tandem repeat benchmark dataset (GIABTR) v1.0^49^ and Truvari^50^. We assessed two catalogs that are available in DRAGEN that differ in the number of STR loci analyzed. The first catalog consists of 50,069 regions where the F-measure (19.68%) was largely driven by the small size of the catalog compared to the 1.7 million regions represented in GIABTR, which impacts the recall. Nevertheless, the precision was high at 95.47%. When utilizing the larger STR catalogs available in DRAGEN which include 174,300 regions, the F-measure improved to 55.13% with the same precision. To provide context to these results, we benchmarked another short-read caller, GangSTR^51^, and compared its performance to DRAGEN’s. Since GangSTR is optimized for a different set of 832,380 regions, we evaluated performance on the intersection of both methods’ analyzed regions against GIABTR (∼174K regions, see **methods**). When restricting the benchmark to the intersection between the two catalogs, DRAGEN achieved a better F-measure of 96.72% (vs 69.86% by GangSTR). When we extended the benchmark to cover all GIABTR regions, DRAGEN’s F-measure for ∼50K and ∼174K catalogs was 94.56% and 94.47%, respectively, whereas GangSTR achieved a F-measure of 62.55% (Figure 2G**, Supplementary Table S5**).

There are two pharmacogenomics related methods that assess *CYP2D6* and *CYP2B6* alleles. For HG002, the caller reported a *1/*U1;*2/*5 star alleles for *CYP2B6* and *2/*4 for *CYP2D6*. The *1 and *U1 alleles in the first genotype represent the reference allele and specific variant in the gene that has reduced enzyme activity, respectively. Similarly, the second genotype, *2/*5, indicates the HG002 sample may carry two different variants of the *CYP2B6* gene which reduces the enzyme activities of the gene. The *CYP2D6* caller for HG002 generated *2/*4 star alleles which indicate the sample carries two haplotype variants that are also associated with enzyme reduction of the gene. The methods for *HBA* 1/2 (Alpha-thalassemia) reported no detected target variants. DRAGEN *HLA* typing on sample HG002 reveals A*01:01, A*26:01, B*35:08, B*38:01, C*04:01, and C*12:03 class I alleles and DQA1*01:05, DQA1*03:01, DQB1*03:02, DQB1*05:01, DRB1*10:01, and DRB1*04:03 class II alleles. Class I genotyping results are concordant with HLA-LA^53^, another WGS based HLA caller. For *SMN* caller, HG002 has “negative” affected status and carrier status, zero copy numbers of *SMN2Δ7–8* (deletion of exon 7 and 8), 3.77 estimated total copy number i.e., indicates four haplotypes across the two genes. DRAGEN also includes two methods that have been previously published. One method assesses *GBA* and *GBAP1* interactions that can be important for neurological diseases^28^ and reports whether the sample is bi-allelic or not, the total copy number, the carrier status, etc.. For the HG002 sample, DRAGEN reported four total copy numbers and False for both “is_bi-allelic” and “is_carrier” fields. The other method assesses the *LPA* copy number status which provides important information on cardiovascular disease risk^29^. Interestingly this method provides phasing information for ∼50% of the samples. HG002 has 39 *KIV-2 LPA* repeats with allele specific (allele1 and allele2) copy numbers of 25 and 14. These methods are highly specialized for their individual targeted regions of the genome and report important allelic information rather than variants (e.g., single SNV). **Supplementary Table S6** contains the descriptions about callers and results for the HG002 sample.

Since STR, SV, and CNV calls each cover a broad range of variant lengths, it is possible for a single variant to be present in more than one result. Therefore, we developed a procedure to combine DRAGEN STR, SV, and CNV calls together to form a comprehensive deduplicated large variant VCF file using Truvari^50^. The merge procedure analyzed a total of 55,414 variants for HG002 and identified 993 redundant variant representations. To establish the accuracy of the merging, the variants that are labeled SV were extracted from the merged file and benchmarking performed using GIAB SV (v0.6) callset. The benchmarking results of the original SV calls were compared with the benchmarking results after merging and found to be nearly identical with only 36 variant representations altered enough to change their benchmarking status (**Supplementary Figure S1)**.

Benchmarking the DRAGEN pipeline shows it produces accurate results that improve variant performance across all variant types and lengths. The pipeline generates the first fully comprehensive representation of a human genome including all variant types at scale and cost.

### DRAGEN improves the identification of variants across human populations

With the performance of DRAGEN on HG002 characterized, we next applied the pipeline to other standard GIAB reference samples to assess the accuracy and comprehensiveness of DRAGEN across multiple ethnicities. These samples include the HG001 (NA12878) sample, the parent samples of AshkenazimTrio (HG003 and HG004) and the ChineseTrio samples (HG005, HG006 and HG007). Figure 3A shows an overview of results across variant types and size regimes. An average of 4,934,765 SNV were detected per sample (substitutions: 3,987,380, small insertions: 461,743, and small deletions: 463,072). A balance (ratio: 0.999) between small insertions and deletions was observed. The mean SNV transition/transversion ratio was observed to be 1.98 and total HET/HOM ratio to be 1.49. For structural variants (SV: >=50bp), the mean SV count per sample was 14,734 with a range between 14,093 and 15,224 per individual. Across samples insertions (mean: 48.78%) were the most frequently occurring SV type, followed by deletions (mean: 39.10%), translocations (mean: 5.20%), inversions (mean: 1.37%), and tandem duplications (mean: 0.36%) (**Supplementary Table S7**). This follows the expected distributions of insertions being the most frequent variant type, which is typically not observed by other Illumina based methods^7^. DRAGEN calls other variants such as copy numbers, short-tandem repeats, as well as variants for some complex and medically relevant genes. On average, 632 CNVs per sample (range between 583 and 718) were detected with lengths between 1kbp and 500kbp (**Supplementary Table S7**). The STRs were detected across 50,069 loci including 62 known pathogenic loci for each sample. Across the samples an average of 13,690 heterozygous and 8,901 homozygous STR variant calls were identified.

**Figure 3:**
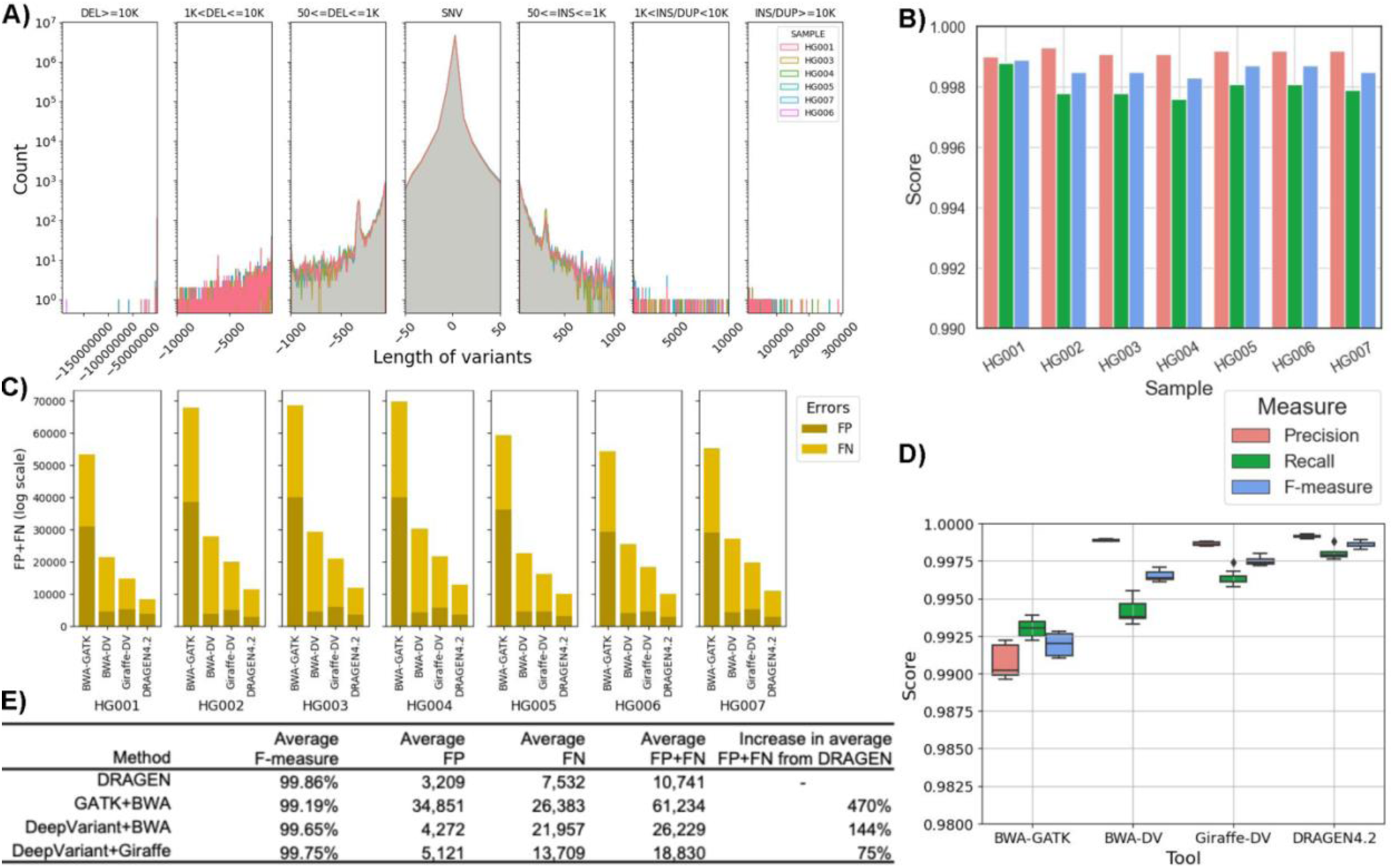
Performance overview of DRAGEN for HG001-07 samples. **A)** Length distribution of different variants for all samples (bin sizes used for the plot from left to right are: 500, 250,150,50300,150, 250, 500); **B)** The recall, precision, and F-measure of DRAGEN for HG001-07 samples; **C)** The comparison of False negative (FN) and False positive numbers among DeepVariant with BWA, DeepVariant with Giraffe, and DRAGEN for HG001-07 SNV calls; and **D)** Comparison of recall, precision and F-measure of these samples for four different tools i.e., DRAGEN, GATK, and DeepVariant-BWA, DeepVariant-Giraffe. **E)** The average F-measures, and errors (false positives and false negatives) for different tools.

DRAGEN performance was then evaluated against the GIAB v4.2.1 benchmarks for HG001-7 samples for SNV and indel^40^. The recall for genome-wide calls were in the range of 99.96% and 99.88% with precision between 99.90% and 99.93% (**Supplementary Table S7**). For SNV and indel, the mean F-measures were 99.87% and 99.79%, respectively (Figure 3B). This shows a remarkably high consistency across all samples in the performance to identify SNV and indel. DRAGEN SNV call performance was then compared against GATK and DeepVariant (DV) calls with BWA and Giraffe^15^ mapper using the GIAB benchmark for all these samples (Figure 3C and D and also see **methods**). Across all callers and samples, the F-measure was below DRAGEN’s: GATK 99.10% to 99.28%; DV-BWA: 99.61% to 99.71%. The higher F-measure is largely attributable to improved detection of substitutions and indels (**Supplementary Table S7**). The benchmarking across all seven samples (HG001-07) allows further assessment of the ability of DRAGEN to utilize a multi genome (graph) reference. Figure 3C shows the accuracy of DRAGEN compared to the accuracy obtained by aligning on the HPRC reference pangenome with Giraffe^15^ and variant calling with DeepVariant (DV) ^45^, the BWA ^54^ with DV pipeline, and the GATK pipeline. When compared to GATK+BWA, DRAGEN shows an average error reduction of 82.45% on combined SNV and indel, with an average reduction of 83.49% on SNV and 75.91% on indel. When compared to DeepVariant+BWA, DRAGEN shows an average error reduction of 59.06% on combined SNV and indel, with an average reduction of 61.31% and 45.87% on SNV and indels, respectively, confirming the trend observed in the previously reported precisionFDA V2 samples^55^. When compared to Giraffe-DV, DRAGEN reports an average error reduction of 42.91% on combined SNV and indel, with an average of 44.00% on SNV and 38.52% on indel.

Since these samples are trios (Ashkenazi (HG002, HG003, HG004), Chinese (HG005, HG006, and HG007)), the variant calling was further validated based on Mendelian inconsistencies. The percent of genotypes at which a trio had “no missingness” and “no Mendelian error” was found to be 97.70% and 96.58% for Ashkenazim trio and Chinese trio, respectively when genome wide analysis was performed. The genotypes’ assigned by DRAGEN were found to have Mendelian error at 2.30% and 3.42% for Ashkenazim trio and Chinese trio, respectively. Nevertheless, when considering GIAB high-confidence regions that encompass approximately 90% of the genome and exclude certain complex segmental duplications and centromere regions, the error rates decreased to 0.15% and 0.33% for the respective trios. The observed De Novo variant rate for both trios were found to be 0.05% (see **methods** and **Supplementary Table S7**).

### Comprehensive variant detection at population scale analysis using DRAGEN

We next applied DRAGEN to discover variants in the well-studied high coverage 1000 genome project (1kGP) ^22,24^ samples and analyze the catalog of genomic variation at population and cohort levels. The 1kGP samples consist of a total of 3,202 samples from 26 different populations of five different ancestry (i.e., super-population) groups: African (AFR), European (EUR), South Asian (SAS), East Asian (EAS), and American (AMR). This collection of samples contains 1,598 (49.91%) males and 1,604 (50.09%) females. The AFR samples have the highest number of samples (n=893, 27.89%), followed by EUR (n=633, 21.64%), EAS (n=601, 18.77%), SAS (n=585, 18.27%), and AMR (n=490, 15.30%). Recently, the low-coverage (7.4x) WGS datasets^22^ of 2,504 samples in 1kGP has been expanded to 3,202 high-coverage (35x) dataset^56^. We analyzed the 1kGP with DRAGEN in order to compare with the recently published SNV callset^56^ with GATK and SV callset with a combination of three tools (GATK-SV^57^, svtools^58^, Absinthe^59^). The analysis with DRAGEN showed an improved performance of variant callings in terms of novel small as well as structural variants ^56^.

For this analysis, it is important to have accurate single sample calling methodologies but also to have methods that combine VCF files from multiple individuals and be able to annotate the variants rapidly and accurately. To accomplish this, a new gVCF merge method for SNV was implemented (see methods) and we utilized Truvari to combine STR, SV, and CNV together. This results in two population merged VCF files, one for SNV and indel and one for larger variant classes.

For small variants (<50bp), the DRAGEN Iterative gVCF Genotyper (IGG) can efficiently aggregate hundreds of thousands to millions of gVCFs to perform joint calling and genotyping. This generates a fully genotyped population VCF file, which is needed for any genome wide association studies, rare variant studies, phasing and imputation, and ancestry studies. The output population VCF file also contains cohort level variant statistics (including allele frequency, sample genotype rate, and coverage rate) and QC metrics (such as Hardy Weinberg test p-value and inbreeding coefficient) that can be used for downstream variant filtering (See **methods** for details). Prior to the aggregation, variants with DRAGEN machine learning quality score below threshold QUAL=3 are filtered. The joint call set has an average per sample SNV recall of 99.92%, precision of 99.78%, and F1-measure of 99.85%, and indel recall of 99.84%, precision of 99.71%, and F1-measure of 99.77%, as evaluated based on GIAB samples. The aggregation over 3,202 samples took almost 2 hours on Illumina phase4 server with concurrency of 200 jobs.

For STR, SV and CNV, the variants were first combined on a per individual level to remove redundant variant representations across types using Truvari^50^. Truvari compares the alleles and sizes together with the location and the type of variant event (e.g., deletion vs. insertions). **Supplementary Figure S1** shows this across HG002 with remarkably similar performance values on SV only and merged STR, SV, and CNV results. After this first step per individual, individuals at population level were merged.

#### Population level SNV and indel identification across 3,202 individuals

We applied DRAGEN across 3,202 high coverage (35x) 3,202 1kGP samples to perform the comprehensive variant calls (SNV, indel, SV, STR, CNV) to demonstrate the scalability. The variants were analyzed, and the results were compared with the published results^56^. At cohort level, DRAGEN identified 118,210,374 SNVs and 25,161,418 indel. The Principal Component Analysis (PCA) plot (Figure 4A) for the small variants at the cohort level shows distinct clusters for different populations, which demonstrates shared genetic ancestry among samples. The distribution of SNVs and indel at population level shows that the AFR super-population has the highest number of SNVs and indel (Figure 4B **& C**), due to the higher diversity of AFR but also likely impacted by the high number of AFR samples in the cohort (**Supplementary Table S8**). The average SNVs per sample ranged from 3,762,359 (EAS) to 4,640,044 (AFR) and followed expected diversity^56^. The number of small insertions (<50bp) for EAS (601,678) was lowest and for AFR (833,407) was the highest. This was interestingly inverted when the small deletions (<50bp) were assessed. The highest proportion of singletons (28.7%) was observed in the AFR population, which also follows previous findings. However, EAS has the highest mean singletons (i.e., ratio of total singleton for a population and number of samples) compared to other populations.

**Figure 4:**
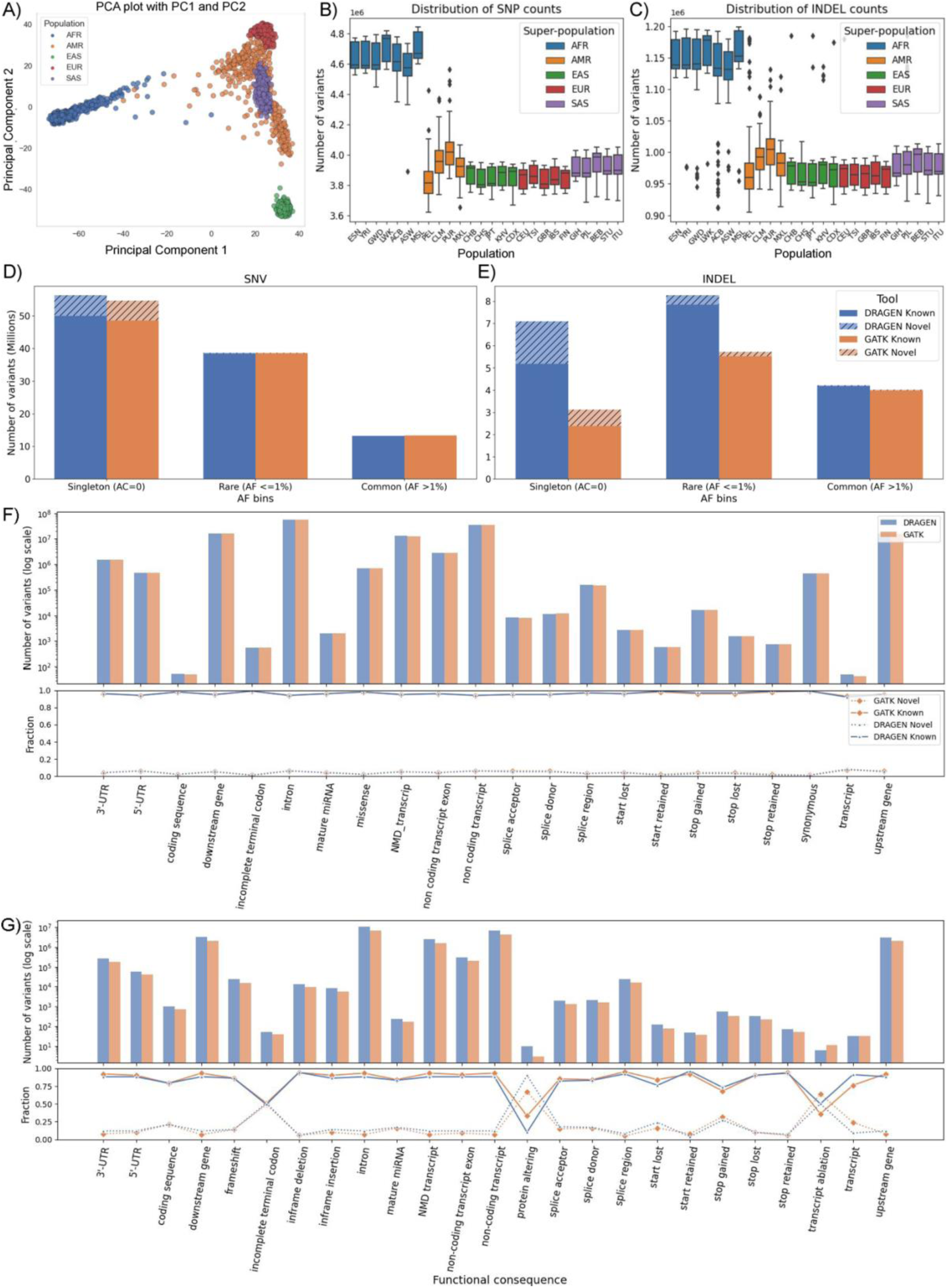
DRAGEN SNV calls for 1kGP sample: **A)** PCA plot of principal component 1 and 2 for SNVs across the population **B)** Distribution of SNV counts and **C)** Distribution of indel counts at super-population level **D)** Singleton (allele count=1), rare (allele frequency <= 1%) and common variant (allele frequency > 1%) counts of GATK v4.1 and DRAGEN v4.0.3 callsets of SNV and **E)** indel across the cohort level. The Known and Novel variants based on dbSNP 155 database **F)** The distribution of SNVs based on their functional annotations shown in the upper plot and the lower plot showing the fraction of Known and Novel variants **G)** The distribution of small insertion and deletions based on their functional annotations.

The allele frequency based analysis on 2,504 unrelated samples shows that DRAGEN generated 56,327,924 (52.03%) singleton, 38,676,117 (35.73%) rare (AF <= 1%), and 13,246,064 (12,24%) common (AF > 1%) SNVs. As compared to previous GATK callset on these samples, it generated 2.95% more singletons and also slightly more rare and common SNVs (Figure 4D-E). For indel variants, DRAGEN generated 7,103,047 singleton, 8,272,115 rare, and 4,226,537 common indel while GATK callset had ∼56% fewer singleton (3,129,240), ∼31% fewer rare (5,727,021), and slightly lower common indel variants (4,023,422). Using the Illumina Connected Annotations (ICA) ^60^ pipeline (also see **methods**), the variants detected by both DRAGEN and GATK callsets were compared with known SNV (dbSNP build 155) to determine which variants were previously observed (i.e. known) or novel. The majority of SNV (93.98%) from DRAGEN were known and 87.86% of indel were known variants. The singleton rate of known variants was ∼50% of SNV and ∼30% of indels (**Supplementary Table S9**).

While most SNV and indel were rare, the novel rate of indel with functional impact was between 9%-15% across samples, while the SNVs novel rate was between 1%-3%. Specifically, among SNVs with functional impact, DRAGEN called 712,630 missense SNVs (94% rare, 2% novel), 441,434 synonymous SNVs (89% rare, 1% novel), and 62,273 SNVs with higher functional impact, including stop/start-gain/lost and splice mutations, (92% rare, 3% novel). For indel with functional impact, DRAGEN called 24,649 frameshift indel (95% rare, 15% novel), 13,185 in-frame indel (91% rare, 9% novel), and 7,707 indel with higher function impact, including stop/start-gain/lost and splice mutations, (81% rare, 9% novel) (Figure 4F-G; **Supplementary Table S10**). We compared the functional annotations of the DRAGEN call set with that of the GATK call set (Figure 4F-G). In the intronic, intergenic and regulatory regions, more SNVs and indels were called by DRAGEN than by GATK. In these annotation categories, the percentage of rare and novel variants (in particular indels) was higher in DRAGEN than in GATK. In annotation categories with low to high functional impact, DRAGEN called fewer missense, synonymous, and functional impact SNVs. The percentage of rare SNVs was higher and the novel SNVs was lower in the DRAGEN call set. Frameshift and functional impacting indels were higher in DRAGEN and found to have a lower allele frequency than GATK. The novel rate was similar between the two call sets, but varied between categories, due to overall lower number of indel in these categories.

The larger number of singletons and novel small variants (<50bp) could highlight DRAGEN’s increased ability to assess repetitive regions of the genome, which is enabled due to the multigenome (graph) implementation (see methods). To answer this, we first focused on the challenging medically relevant gene (CMRG) gene regions that are important for clinical analysis. We analyzed the variants identified by DRAGEN in 389 challenging gene regions and compared them to the previous GATK based results. DRAGEN identified 1,134,340 (0.79% of total) variants in those regions. This is similar to the GATK results of 1,146,580. Next, we investigated if DRAGEN accurately captures the variants in 12 medically important genes that are ill represented on GRCh38^24^. These 12 genes comprise nine which are wrongly duplicated and three that are wrongly collapsed (e.g., 2 instead of 3 copies). These regions include the genes *KMT2C*, *H19, MAP2K3*, *KCNJ18*, *KCNE1, CBS, U2AF1, CRYAA, TRAPPC10, DNMT3L*, *DGCR6 and PRODH*. For the nine genes that are wrongly duplicated, DRAGEN was able to circumvent this bias and reported 35,504 variants. This is in stark contrast to the GATK call set which reports almost 30% fewer (24,249) variants. As an example, for *CBS*, related to cystathionine beta-synthase deficiency^61^, only 221 variants were reported across 1kGP in the previous study^56^ . DRAGEN reported 1,297 variants in its’ call set due to the use of multigenome (graph). For the *H19* gene, related to skeletal muscle disease)^62^, DRAGEN found 341 variants, but GATK found no variants. For genes that were impacted due to collapsed errors, we expected an inflated number of variants due to multiple haplotypes collapsing on top of each other^24^. For these three genes, we observed fewer (20,047) variants from DRAGEN than GATK (24,322). For *MAP2K3,* related to skin and liver diseases^63^, and *KCNJ18,*related to some rare disease^64^, GATK discovered 1,631 more variants than DRAGEN, which are likely false positives^24^ (**Supplementary Table S11**).

#### Unification of large alleles across 3,202 individuals

Next, we investigated the larger variants identified by DRAGEN encompassing STR (50,069 regions), SV and CNV. As described above we merged all large variant types across the samples into one population VCF file. We identified 410,366 STRs (243,778 expansions i.e., reference has fewer repeat units and 166,588 contractions i.e., reference has more repeat units), 1,353,805 SVs (with 262,712 DEL, 620,530 INS, 15,087 tandem DUP), and 6,422 CNVs (3,471 DEL and 2,951 DUP) across the entire 1kGP data set (**Supplementary Table S12**). We first performed a PCA analysis to investigate if these calls follow the expected population structure. Figure 5A shows the PCA colored by super populations. Overall, we observe a nice separation following the population structure in PCA 1 & 2. The large variant PCA has a highly similar structure to that observed in the small variant PCA. The stratification is likely also driven by the higher variant numbers we observe across the African population compared to the other ethnicities, which is also similar structure that was observed in small variant PCA. Figure 5B and C shows the distribution of insertions and deletions per population. Across all SV types we see the expected distributions of variant counts with a slight increase of insertions over deletions. While it remains challenging to identify insertions from short reads, we see the relatively high numbers of DRAGEN insertions obtained following the general population structure. Figure 5D shows the average number of SV per individual for each population. Interestingly, while we observe increases of insertions and deletions for Africans compared to other populations, the same is not observed on duplications or inversions.

**Figure 5:**
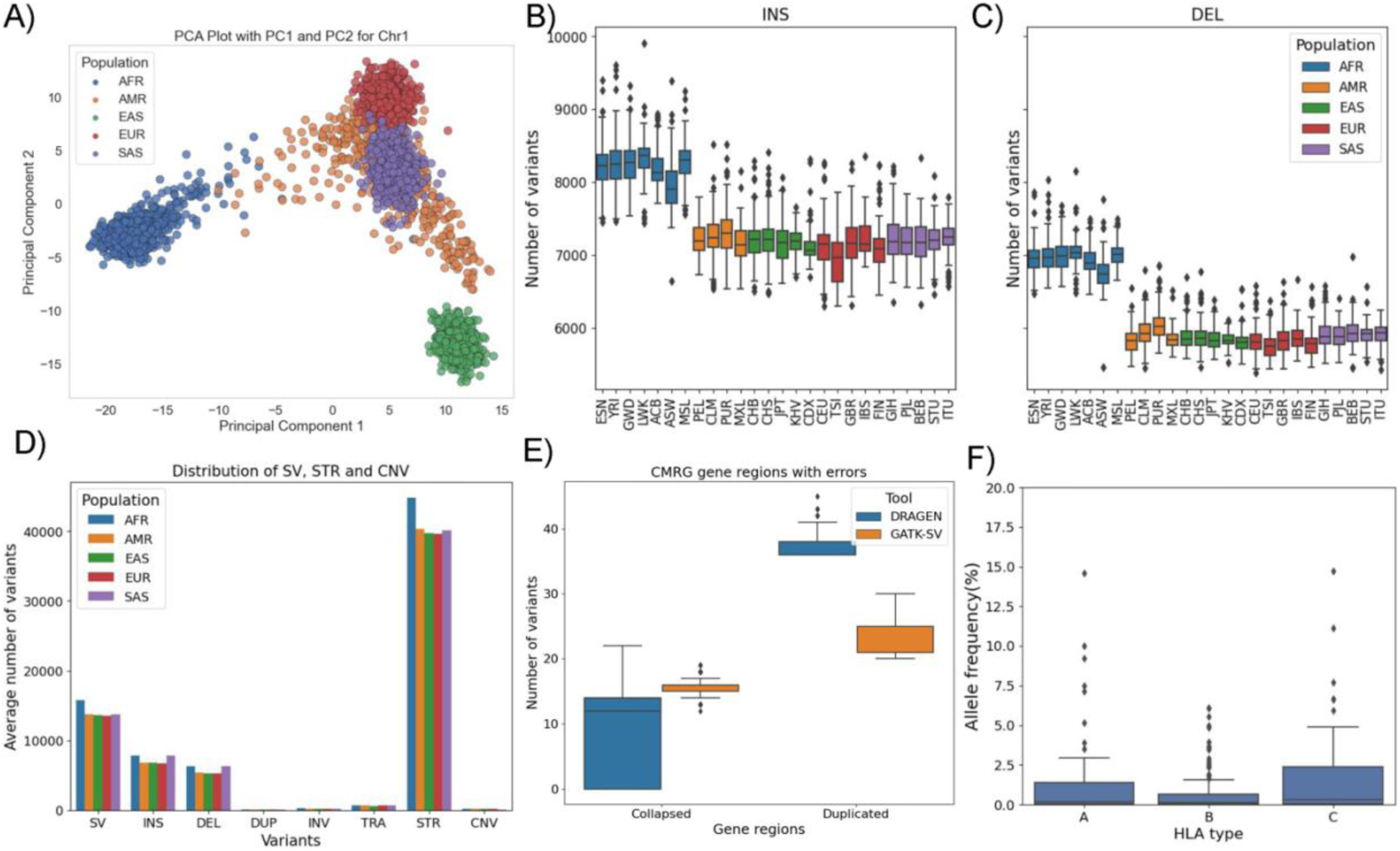
DRAGEN SV calls for 1kGP sample: **A)** PCA of merged STR, SV and CNV for deletions >5% on chromosome. **B)** Distribution of insertion and **C)** deletion type structural variants (>= 50bp) among populations **D)** Distribution of SV, STR (for ∼50K loci) and CNV variants based on average count i.e., total variants of a population / population count **E)** distribution of variant numbers among all 3,202 samples for the 12 challenging medically relevant gene (CMRG) regions (in GRCh38) that are impacted due to falsely duplication and falsely collapsed errors. DRAGEN uses the corrected reference as a part of its multi genome approach to correctly identify more variants in duplicated and in collapsed regions. **F)** Class I HLA allele frequency distributions among all 1kGP populations

We next investigated how many of these variants have been identified previously^19,56^. For this task we used ICA to annotate variant intersections to 1kGP, gnomAD and TOPMed. Across all variants we observed 1,410,769 known variants and 12,459,468 novel variants. **Supplementary Table S13**. contains the distribution based on allele frequency. To cross check consistency of the data set we correlated the allele frequency of the call sets for the overlapping variant calls. We observed a positive correlation (Pearson correlation coefficient: 0.999, p-value=0.0) between allele frequency and the count of variants from the 1kGP database. Next, we checked for the overlap of variants with exonic, intronic and intergenic regions. A total of 65,025 SV (24,464 DEL and 14,395 INS) overlapped exonic regions, 717,559 overlapped intronic regions (348,594 INS and 146,161 DEL), and 602,762 overlapped intergenic regions (257,847 DEL and 116,349 DEL).

Since DRAGEN identifies multiple categories of large variants (SV, STR, CNV) we queried the total number of base pairs impacted across the genome by these variants compared to the small variants. The individual number of variants fluctuate per category - however, categories with fewer variants often contain longer variant alleles. The average number of basepairs impacted by SNV and indel per individual across the 1kGP cohort was 8,618,694 bp while for STR, SV and CNV an average of 8,555,084 (6,427,623 for SV, 860,424 for CNV and 1,267,036 for STR) bp. The number of impacted base pairs by large variants (STR, SV, CNV) is observed to be very close to the number of basepairs impacted by SNV. However, the DRAGEN callset for SV also include many insertion variants (average ∼25% per sample) for which the length of variant was not resolved. Therefore, including these large insertion variants could substantially increase the number of impacted base pairs. This confirms previous reports that the impacted bp from SV are higher than SNV and indel alone and underlines the importance of STR, SV and CNV^7,65^. The AMR population (62,294bp) again showed the highest average bp changes (**Supplementary Table S14**).

We further obtained insight into the SV diversity along the medically important genes. As the 1kGP samples represent healthy individuals, their SVs could be used as controls in studies aiming to identify potentially pathogenic variants. We compared DRAGEN SVs to results that were recently published^56^ from a joint calling ensemble approach (GATK-SV^57^, svtools^58^, Absinthe^59^). Across the 5,030 challenging medically relevant gene regions DRAGEN identified 265,317 variants (197,191 SV; 66,446 STR; 18,038 CNV). The SV callset that was published in the recent studies reported only 27,166 SV with 8,093 insertions and 13,506 deletions. These can be split in mainly 27,884 more deletions and 87,639 more insertions that are discovered by DRAGEN. Within these medically relevant genes there are 12 genes that often suffer in the analysis due to reference biases^24^. As mentioned before, some genes suffer from wrongly collapsed copies which leads to an increased number of variants^24^. On the other side there are several genes that have been wrongly reported multiple times across the genome which often led to missing variant calls due to their repetitiveness^24^. For the duplicated and collapsed regions, a total of 65 and 384 large variants were identified by DRAGEN and the majority of them are SV (95.38% and 97.14%). In contrast, the previous study only reported 36 SV in collapsed and 19 SV in duplicated regions across the entire 1kGP. At the cohort level, on an average each individual has 11 variants that were identified in the erroneous regions. For the AFR population, the average number of variants was 13 and for other populations it was between 9 to 10 variants per sample. The distribution of total number of variants by DRAGEN at the duplicated erroneous regions are higher than the numbers reported in the previous studies and the numbers are lower in the collapsed regions. This shows the improvement of variant calling by DRAGEN that incorporates the corrected regions during variant calls (**Figure 5E** and **Supplementary Table S15**). A lower number of variants is expected in the collapsed erroneous regions if the corrected reference is used as these erroneous regions in the original GRCh38 reference with more than one copies are collapsed into one.

#### Insights across medically relevant but complex genes across 3,202 individuals

Lastly, we investigated results from the DRAGEN specialized gene callers (e.g., *CYPB26*, *CYPD26*^26^, *GBA*^28^, *HLA*, *SMN1&2*^27^) to obtain deeper insights into potential pre conditions across the 1kGP data set. Furthermore, this data set can be leveraged as population controls for these important but complex genes.

For CYP2B6 caller, 2,017 samples had genotypes containing two haplotype specific star alleles (filter status PASS), 1,174 samples had more than one possible genotype and 11 samples (10 AFR and 1 EUR) had no calls reported. The metabolizer status reported in these calls shows that among samples with PASS filter 1,189 with normal metabolizer, 381 with poor, 154 with rapid, 7 with ultra-rapid, 224 with intermediate and 57 with indeterminate status and 858 samples are with *1/*1 genotype. Among the samples with multiple genotypes, 945 of them are with genotype “*1/*6;*4/*9”. For CYP2D6 calls across all samples, only two samples (one EUR and one SAS) had more than one possible genotype. There were 11 with no calls (2 AFR, 1 SAS, 6 EAS and 3 AMR) and the remaining 3,188 samples had one genotype with two haplotype specific star alleles. The metabolizer status showed that 1,557 samples had normal status, 733 intermediate status, 59 poor, 106 ultrarapid and 143 indeterminate status.

For GBA^28^ caller that detects both recombinant-like and non-recombinant-like variants in the GBA gene, DRAGEN reported no samples with any presence of a recombinant-like variant on each chromosome (homozygous variant or compound heterozygous). However, it reported 13 samples (3 AFR, 5, EUR, 1 AMR, 1 SAS and 3 EAS) with presence of a recombinant-like variant on only one chromosome. The reported total copy number values showed that the majority of samples (95.47%) had aggregate CN of 4 across gene and pseudogene. Only 16 samples had an aggregate CN of 3, and the remaining 129 samples (111 AFR, 1 EUR, 6 AMR, 11 EAS and 15 SAS) had aggregate CN in the range between 5 and 10. It reported only one sample (of EAS) that has one deletion breakpoint in GBA gene which indicates if the sample has one of the recombinant-like deletion variants.

For SMN caller, DRAGEN reported SMA affected status as “false” for all samples and SMA carrier status “true” for 49 (1.53%) samples (3 AFR, 19 EUR, 12 AMR, 7 EAS and 8 SAS). This is in the range of rates of carriers, which is between 2.50%-1.67% across the population ^66^.The copy number of SMN1 was reported to be 2 for majority of samples (2,428) and it was not reported for 19 samples (None for SMN1_CN). For SMN2 copy number, 395 samples with 0 CN, 1,275 with 1,427 with CN 2, 86 with 3 or 4 and 19 with no reported copy number.

DRAGEN HLA caller reports HLA typing results of six class I alleles (i.e., A-1, A-2, B-1, B-2, C-1, C-2), it was reported 60 distinct alleles for A-1, 70 for A2, 121 for B-1, 132 for B-2, 43 for C-1 and 57 for C-2. For A-1 type, A*02:02 was reported to be allele with highest allele frequency of 15.8% that followed by A*11:01 with 11.62% and the remaining alleles were within 0.03% and 10.06%. For A-2 type, the allele A*02:01 was reported to be with highest allele frequency of 13.34% and all others were within 0.03% and 9.90%. For B-1 type, the allele B*07:02 was with highest allele frequency of 6.71% and the remaining alleles were in between 0.03% and 5.78%. For B-2 type, the B*35:01 allele had highest allele frequency of 6.62% and remaining alleles were in between 0.03% and 5.68%. For C-1 type, the highest allele frequency of 17.36% was reported for the allele C*04:01 and the remaining alleles were in the range between 0.03% and 13.46%. Lastly for C-2 type, again the allele C*04:01 reported to have the highest allele frequency of 12.05% and others were within 0.03% and 8.81%. The allele frequency distribution of HLA type-1 classes among all 1kGP populations are shown in **Figure 5F**. **Supplementary Table S16** has details for *HLA* type counts.

Thus, throughout the paper we have demonstrated the accuracy and scalability of the DRAGEN framework across all variant types. We have demonstrated this across all different variant classes across a wide spectrum of human population and with a focus on genome wide as well as medically relevant genes. This revealed many novel variants (SNV - CNV) that were not detectable in previous studies of this data set. Furthermore, we were able to provide this more comprehensive call set together with the results of the specialized callers as a population reference for future studies.

## Discussions

In this paper, we present a novel method DRAGEN to comprehensively identify all germline variants at scale. DRAGEN includes novel methods to improve the identification of SNV, indel, STR, SV, CNV and nine targeted callers, which is further promoted by the utilization of a multi genome (graph). As such it represents the first application that can utilize multigenome (graph) across all types of variants and truly highlights a significant step in the analysis of Illumina sequencing data. Even more impressively, DRAGEN achieves this high accuracy while providing a fast and scalable method that is able to process a 35x whole human genome Illumina fastq within ∼30 min of computation time with an onsite DRAGEN server achieving F-scores from 76.90% (SV) to 99.86% (SNV) across the different variant classes. In addition, we introduce novel methods to compare and merge the different variant classes across population data to obtain fully genotyped VCF files for SNV and indel at high precision. Furthermore, Truvari^50^ can be leveraged to combine STR, SV and CNV together across a set of genomes. Thus, DRAGEN enables the assessment of variants at unprecedented scale and accuracy, which will further enable new insights into medical and biological research. As such DRAGEN is currently already deployed at multiple large-scale projects such as UK Biobank (UKBB), All of Us (AoU) to name only two. This enables comprehensive variant calling but even more comparability across large scale cohorts to leverage each other’s results to improve personalized medicine and research applications. To further promote this DRAGEN is now getting directly integrated into the Illumina sequencing machines. To further promote this DRAGEN is now getting directly integrated into the Illumina sequencing machines.

Over the past decade we and others have highlighted that not only SNV and indel are impacting important phenotypes (e.g. medical) but also SV and CNV are more and more often reported for certain diseases^67,68^ such as cancer, rare genetic disorders etc. Furthermore, STR mutations are often reported with pathogenic alleles (e.g., FMR1) that impact adult diseases such as neurological disease and many more^49,69^. In addition, current disease research is often focused on rare diseases that require a significant amount of probands and controls to decipher statistically significant signals of mutations impacting certain genes or pathways leading to a certain disease phenotype. Thus, it is of utmost importance to promote the identification of all variant types (independent of size and complexity) at scale across thousands or millions of samples. We showcased the speed and scalability across multiple human populations. Nevertheless, variant identification especially for STR and SV remains challenging for short reads. This is due to repeats and the complexity of these alleles^7^. Despite those challenges, we demonstrate a significant improvement of SV, CNV and STR discovery compared to other state of the art methods. This highlights that while the signals of the alleles are present even for complex alleles in short reads, it requires advanced approaches to decipher and report them accurately.

This is in part enabled by leveraging multigenome (graph). This version of DRAGEN includes 64 haplotypes that represent human populations well. Others will be added over the time as they become available. Using the current set of 64 haplotypes, DRAGEN outperforms existing pan genome implementation (e.g. Giraffe^14^) not only in accuracy (SNV: 99.85% vs 99.74% F-score) but further in scalability and runtime. The advantage of the graph by including multiple haplotypes is the better representation of common variants (here AF > 1%). In addition, the DRAGEN multigenome (graph) is already incorporated for SV and CNV calling, something that is not possible right now with any other graph genome implementations since they focus primarily on genotyping variants ^14,70^. DRAGEN analyzes variants using the multigenome (graph) with the variant coordinates projected back to either GRCh38 or CHM13 (not shown here). To further promote the scalability of the method at population level we have presented new approaches to provide population level VCF files that are required for any subsequent GWAS or otherwise functional studies. Here we presented IGG to obtain a fully genotyped multi-sample VCF file. We demonstrated that we identified many novel variants not only genome wide but also in important medically relevant genes. Furthermore, we overcame the challenge of combining STR, SV and CNV variants at an individual and population level. This is now implemented over Truvari, which first merges across variants within an individual and subsequently across individuals. We have evaluated both merging strategies in this manuscript. This allows more comprehensive insights per sample and will foster new findings across population studies across different phenotypes. For the 1kGP cohort dataset, DRAGEN was able to discover more variants i.e., SNV, indel (2-50bp) and large variants (>=50bp) as compared to the recently published results on the same cohort. Besides these variants, DRAGEN also discovered short tandem repeat expansions for ∼50K loci and the copy number variations (>=1kbp) across the genomes. Still certain genes/regions of the genome require special attention (e.g., *HLA*, *CYP2D6*, *CYP2B6*, *LPA* etc.). For this, DRAGEN includes specialized callers that resolve genes (e.g., *SMN1*, *LPA*) that are of high importance but often escape genome wide analysis. These nine specialized callers have now been all integrated in the same platform, again promoting the notion of the most comprehensive genome analysis to date.

Thus overall, DRAGEN represents a significant milestone in the analysis of sequencing data and will lead to novel insights across many diseases from Mendelian over rare diseases being the only platform that is highly comprehensive but also scalable.

## Methods

### DRAGEN Overview

DRAGEN (Dynamic Read Analysis for GENomics) is a bioinformatics platform developed by Illumina that is designed to accelerate and improve the analysis of genomic sequencing data. DRAGEN uses field-programmable gate array (FPGA) technology to accelerate sequence alignment, variant calling, and other computationally intensive tasks that are commonly performed in the analysis of genomic data.

DRAGEN supports a wide range of applications, including whole genome sequencing, exome sequencing, RNA sequencing, oncology, and more. The platform is designed to be highly scalable, allowing it to process large amounts of data quickly and efficiently, and it is optimized for use in high-throughput sequencing environments. While DRAGEN can be used in a wide range of applications, including cancer research, infectious disease studies, and population genetics, here we focus on demonstrating its capabilities in the whole genome sequencing (WGS) germline context.

DRAGEN’s capabilities for whole genome germline applications include 1) Fast end to end analysis due to FPGA hardware acceleration 2) Comprehensive variant calling: DRAGEN includes methods to detect a wide array of variant types under a single command line, such as single nucleotide variant (SNV) and insertions/deletions (indel), structural variants (SV), copy number variants (CNV), short tandem repeat expansions (STR), targeted callers to detect pathogenic variants and/or gene conversion events in challenging medically relevant genes (CMRG), and joint*/de novo* variant calling. 3) Scalability: DRAGEN is designed to be highly scalable, meaning it can process large amounts of data quickly and efficiently. This is particularly important for WGS applications, large cohorts analysis for population genomics studies. 4) Streamlined workflow: DRAGEN offers a complete and automated end-to-end solution to map and align raw sequencing reads and output variants in VCF files, that can then be interpreted downstream.

### DRAGEN Read Mapping

DRAGEN uses hash-table based seed mapping into the reference genome, with dynamic seed extension as needed to reduce *k-mer* match sets to manageable sizes. Rescue scans search the expected insert-length interval for any missing mate matches, and both gapless scoring and gapped Smith-Waterman alignment are used to extend to full-read alignments. Split-alignment possibilities are discovered and scored for each mate, and pair scores are assigned to whole-template alignment candidates, considering the empirical insert length distribution. MAPQ is estimated mainly in proportion to the difference between best and second-best pair scores, separately for each mate. This entire map/align pipeline is executed by FPGA hardware.

For the results presented here, DRAGEN used hg38 reference and hg19 with multigenome (graph) augmentations encoding population haplotype information to improve mapping accuracy. GRCh38 is used here as an example, but the DRAGEN multigenome (graph) can be applied to and constructed for all existing human reference FASTAs (hg19, hs37d5, hg38, chm13). 64 population haplotypes in each genomic region were derived from phased SNV and indel calls for 32 globally distributed samples, with low-confidence variants (under QUAL 30) excluded unless phased with nearby higher-qual ones, and low-AF variants (occurring in fewer than 5% of haplotypes in a larger panel) also excluded. Haplotypes were further restricted to 366Mbp of the most difficult-to-map regions in hg38.

Two types of multigenome (graph) augmentations are derived from these population haplotypes during reference construction. Firstly, isolated SNVs (not phased with other variants within 150bp) are represented as multi-nucleotide IUB codes injected into the hg38 sequences. These multi-base codes have two effects: the mapping hash table contains additional *k-mers* for seed positions overlapping them, and alignment scoring considers multiple read bases to be matches.

Secondly, indels and/or phased clusters of SNVs are represented as alternate sequences (alt contigs) in addition to the hg38 sequences. Each alt contig has a known liftover alignment into hg38, which is critical to alignment treatment during read mapping. Additional seed *k-mers* from alt contigs are populated into the mapping hash table, where they point back to their source alt contig positions but are organized together with corresponding primary-contig *k-mers* to encode their liftover relationship. At each position where an alt-contig *k-mers* differs from its primary-contig liftover image, a copy of this alternate *k-mers* is added pointing to the primary-contig liftover position, improving seed mapping sensitivity in the primary contig.

Reads thus find seed mappings into both primary and alt contigs. The seeds’ liftover relationships are imported from the hash table, organizing scored alignments into “liftover groups”, each typically with one primary-contig member and one or several alt-contig members. Alignment comparison, winner selection, and MAPQ estimation are then performed at the level of liftover groups rather than individual alignments, each liftover group using the highest alignment score among its members. The winning liftover group’s primary-contig member is always the one reported in SAM/BAM output, which facilitates variant calling in hg38 coordinates.

These graph augmentations improve alignment accuracy by enabling reads to effectively achieve better alignment scores at hg38 sites where their differences correspond to variants in the population haplotypes. A particular read may, for example, score equally well in both a gene and its pseudogene as represented in hg38, but if its differences match population haplotypes observed to occur only in the gene, then this read is granted an improved score in the gene, and will map there with positive MAPQ to support calling those variants in the gene.

### DRAGEN Germline Small Variant Caller

The DRAGEN Germline Small Variant Caller is a haplotype-based variant caller that takes mapped, aligned and sorted DNA reads as input, calls single nucleotide variants (SNV) and indels (insertions and deletions), and outputs as a (g)VCF file (**Supplementary Figure S2**). DRAGEN includes a sample-specific characterization step, which takes as input the aligned BAM, and outputs estimates of indel error rates, which then inform the parameters for the Hidden Markov model (HMM) that performs the read likelihood calculation inside the germline variant caller.

The DRAGEN germline variant caller workflow is described in **Supplementary Figure S3**. The first step (step 1 in **Supplementary Figure S3**) looks for sufficient coverage and evidence of variants in the reads to establish active regions. Since DRAGEN is a haplotype-based variant caller, the reads covering an active region are then locally assembled via a *de Bruijn* graph to generate a set of candidate haplotypes (Step 2 in **Supplementary Figure S3)**. This step is similar in concept to GATK4/Mutect2^71^. Once the haplotypes are assembled, they are aligned against the human reference to identify candidate variants. It is possible to augment the events generated by the graph by recruiting events from “column-wise” detection which consists of counting the number of reads supporting a variant at a given column in a read pileup. The HMM then computes a likelihood for each read-haplotype pair, considering the indel sample-specific noise estimates computed upstream of the variant caller (step 3 in **Supplementary Figure S3**). In the genotyper (step 4 in **Supplementary Figure S3**), candidate genotypes are formed from diploid combinations of variant events (SNV or indel).

Given a set of reads 𝑅 = {𝑟_1_. . . 𝑟_𝑁𝑅_ } and a set of haplotypes 𝐻 = {ℎ_1_. . . ℎ_𝐻_}, the HMM produces scores 𝑃(𝑟_𝑖_ |ℎ_𝑘_) for all combinations of 𝑖 = 1. . . 𝑁_𝑅_ and 𝑘 = 1. . . 𝑁_𝐻_ . At a given locus, we have a set of candidate alleles 𝐴 = {𝑎_1_. . . 𝑎_𝑁𝐴_ }. Let ℎ_𝑘_ → 𝑎_𝑗_ indicate that haplotype ℎ_𝑘_ contains allele 𝑎_𝑗_. The goal of genotyping is to calculate the posterior probability 𝑃(𝑎_𝑗_, 𝑎_𝑘_|𝑅), 𝑗 = 1. . . 𝑁_𝐴_, 𝑘 = 1. . . 𝑗 .

For each allele 𝑎_𝑗_ (including the reference allele), the conditional probability 𝑃(𝑟_𝑖_|𝑎_𝑗_) of observing a read 𝑟_𝑖_ given the event 𝑎_𝑗_ is estimated as the maximum 𝑃(𝑟_𝑖_ |ℎ_𝑘_) across all haplotypes supporting the event.

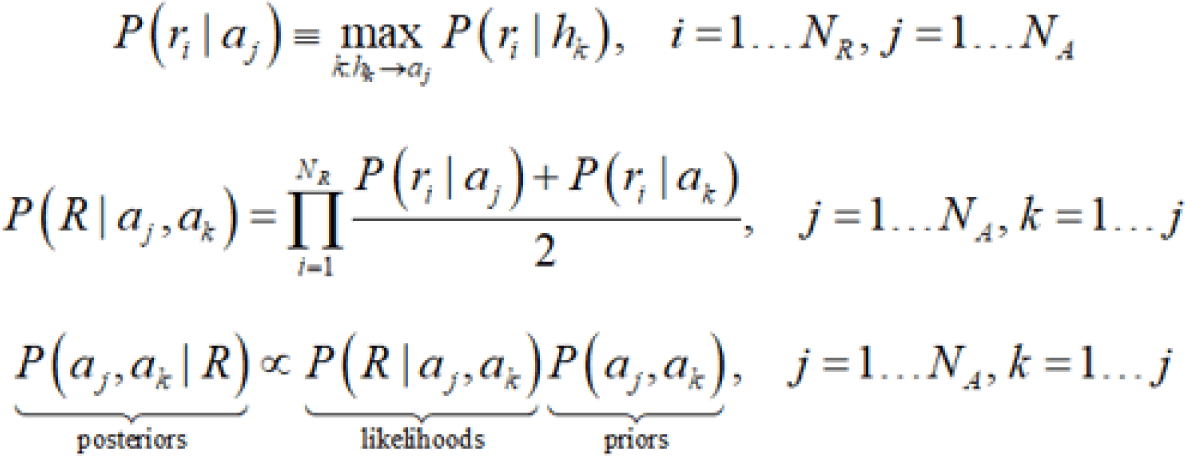

These conditional probabilities 𝑃(𝑟_𝑖_|𝑎_𝑗_) are combined into the conditional probability 𝑃(𝑟_𝑖_ |𝑎_𝑗_, 𝑎_𝑘_) for a genotype (event pair) and multiplied to yield the conditional probability 𝑃(𝑅|𝑎_𝑗_, 𝑎_𝑘_) of observing the whole read pileup. Using Bayes’ formula, the posterior probability 𝑃(𝑎_𝑗_, 𝑎_𝑘_| 𝑅) of each diploid genotype is calculated, and the winner is called (based on the maximum posterior probability). The genotyping matrix is normalized by the sum of all genotypes and the variant quality score (QUAL) is computed as *-10log10* of the posterior probability of the homozygous reference genotype. The QUAL field is updated when machine learning recalibration is enabled. This yields better calibrated QUAL values. Finally, a set of simple hard-filtering rules may be applied to the output VCF to find the best tradeoff between sensitivity and precision (highest F-measure).

#### Key Innovations of the DRAGEN Germline Variant Caller

The germline variant caller incorporates advanced methods which provide substantial improvements in the analytical precision and sensitivity compared to existing third party tools (see **results**). These methods address key variant calling challenges: 1) Sample-specific PCR noise, 2) Correlated pileup errors, 3) Overlapping candidate events, 4) Local assembly failures and Incomplete haplotype candidates.

#### Sample-Specific PCR Error Model

One of the challenges in variant calling is distinguishing indel errors from true variants. To do so, variant callers often employ a Hidden Markov Model (HMM), which models the statistical behavior of indel errors, as part of the probability calculation. The HMM typically has input parameters, Gap Open Penalty (GOP) and Gap Continuation Penalty (GCP), which are directly related to the indel error rate (i.e., indel error rate = f(GOP,GCP)). Indel errors are more likely in the presence of short tandem repeats (STRs), and the error probability (and thus GOP and GCP) may depend on both the period and the length of the STR. The error process may differ significantly from one dataset to another, depending on factors such as PCR amplification. For accurate detection, it is important to use HMM parameters that accurately model the error process on a per sample basis. However, typical variant callers often use fixed parameters or non-sample-specific predetermined functions that fail to accurately model the error process, resulting in poor detection performance.

The HMM auto calibration implemented in DRAGEN addresses the above problem by estimating the PCR parameters directly from the dataset being processed. This operation is performed after mapping & alignment and prior to variant calling, without knowledge of the ground truth and without using external databases of known mutations. The parameters depend on both the STR period and the repeat length.

For a given STR period and length, a set of N loci with the desired period and length is selected, and the algorithm examines the pileups of reads mapped to those loci, counting the indels observed at each locus. The key idea is that by considering a sufficient number of loci, it’s possible to accurately estimate the parameters of interest. We do so by finding the parameters that maximize the probability of producing the set of N observed pileups. If the number of parameters to maximize the probability over is small enough (e.g., 2 or 3), an exhaustive search is possible. In the current implementation of DRAGEN, the optimization is performed over three parameters: GOP, GCP and alpha, where alpha indicates the probability of indel variants of any non-zero length. For each STR period and length considered, the search outputs GOP, GCP and alpha that maximize the probability of producing the set of N observed pileups, and those values are used as input to the HMM.

#### Modeling Sources of Correlated Pileup Errors

##### Foreign Read DETECTION (FRD)

Conventional variant callers treat mapping errors as independent error events per read, ignoring the fact that such errors typically occur in bursts (causing correlated mapping errors). This can result in variant calls emitted with very high confidence scores in spite of low MAPQ and/or skewed AF. To mitigate this problem, conventional variant callers typically filter out reads upstream of variant calling, based on a MAPQ threshold (i.e., reads with MAPQ< threshold are excluded from the calculation). However, this discards valuable evidence from within the variant caller and does a poor job of suppressing false positives. To handle correlated mapping errors, FRD extends the genotyping algorithm by incorporating an additional hypothesis that some read(s) in the pileup are foreign reads (i.e., their true location is elsewhere in the reference genome). The algorithm exploits multiple read pileup properties like relative allele depth, localized reads, MAPQ per read, and strand bias and incorporates this evidence into the probability calculation in a mathematically rigorous manner.

New genotype candidates hypotheses are added to the legacy list of diploid genotypes (those that assume independent pileup errors). For example, in the case of a locus with 1 ALT allele, in addition to considering P(G00|R), P(G01|R), P(G11|R), we add two more hypotheses as P(G00,F1|R) and P(G11,F0|R), where allele F0 and F1 represent reference allele and ALT allele coming from a mapping error. The properties of those errors, such as allele depth and MAPQ are incorporated in the calculation of P(G00,F1|R) and P(G11,F0|R). Then the winning genotype is taken over max (max(P(G00|R), P(G00,F1|R)), P(G01|R), max(P(G11|R), P(G11,F0|R))). Sensitivity is improved from rescuing FN, correcting genotypes and enabling lowering of the MAPQ threshold for incoming reads into the variant caller. Specificity is improved from removing FP and correcting genotypes.

With FRD, DRAGEN variant caller can apply more relaxed MAPQ thresholds when accepting reads for downstream processing. For example, it takes in reads with MAPQ as low as 1, while other conventional callers apply a more stringent MAPQ threshold (typically 20 or higher) to filter out mid-to-low confidence mapped reads. An overly high MAPQ threshold can cause valuable evidence of variants to be lost, hence being able to lower the MAPQ threshold yields increased sensitivity.

##### Base Quality Drop-Off (BQD)

Conventional variant callers are designed with the assumption that sequencing errors are independent across reads; following this assumption, it’s very unlikely that multiple identical errors will occur at a specific locus. However, after analyzing NGS datasets, it was observed that bursts of errors are far more common than would be predicted by the independence assumption, and these bursts can result in lots of false positives.

Fortunately, these errors have distinct characteristics differentiating them from true variants. The BQD (base quality drop off) algorithm implemented in DRAGEN is a detection mechanism that exploits certain properties of those errors (strand bias, localization of the error in the read, low mean base quality, at the locus of interest) and incorporates them into the probability calculation in a simple and robust manner, in the genotyper. New genotype candidates hypotheses are added to the legacy list of diploid genotypes (those that assume independent pileup errors). For example, in the case of a locus with 1 ALT allele, in addition to considering P(G00|R), P(G01|R), P(G11|R), we add two more hypotheses as P(G00,E1|R) and P(G11,E0|R), where allele E0 and E1 represent reference allele and ALT allele coming from a sequencing error. The properties of those errors, such as strand bias, localization of the error in the read and mean base quality are incorporated in the calculation of P(G00,E1|R) and P(G11,E0|R). Then the winning genotype is taken over max (max(P(G00|R), P(G00,E1|R)), P(G01|R), max(P(G11|R), P(G11,E0|R))).

Being able to characterize correlated sequencing errors from within the core of the variant caller results in a significant gain in specificity because a lot of FP calls are removed. It also helps sensitivity by correcting genotype errors.

##### Joint Detection of Overlapping Events

As described earlier, in the genotyper (step 4 in **Supplementary** Figure 2), candidate genotypes are formed from diploid combinations of variant events (SNV or indel).

Given a set of reads 𝑅 = {𝑟_1_. . . 𝑟_𝑁𝑅_ } and a set of haplotypes 𝐻 = {ℎ_1_. . . ℎ_𝐻_}, the HMM produces scores 𝑃(𝑟_𝑖_ |ℎ_𝑘_) for all combinations of 𝑖 = 1. . . 𝑁_𝑅_ and 𝑘 = 1. . . 𝑁_𝐻_ . At a given locus, we have a set of candidate alleles 𝐴 = {𝑎_1_. . . 𝑎_𝑁𝐴_ }. Let ℎ_𝑘_ → 𝑎_𝑗_ indicate that haplotype ℎ_𝑘_ contains allele 𝑎_𝑗_. The goal of genotyping is to calculate the posterior probability 𝑃(𝑎_𝑗_, 𝑎_𝑘_|𝑅), 𝑗 = 1. . . 𝑁_𝐴_, 𝑘 = 1. . . 𝑗 .

For each allele 𝑎_𝑗_ (including the reference allele), the conditional probability 𝑃(𝑟_𝑖_|𝑎_𝑗_) of observing a read 𝑟_𝑖_ given the event 𝑎_𝑗_ is estimated as the maximum 𝑃(𝑟_𝑖_ |ℎ_𝑘_) across all haplotypes supporting the event.

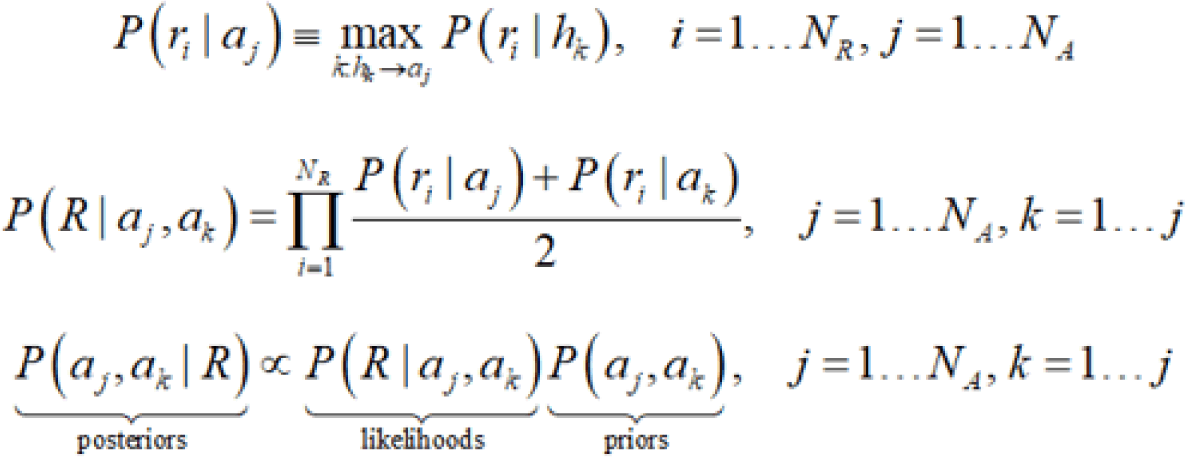

However, the maximum operation over all haplotypes supporting the event 𝑎_𝑗_ is sub-optimal and gives inaccurate variant calls for cases of overlapping events or events separated by a short tandem repeat (STR) region. The optimal solution is to jointly call variants in each region instead of treating each event as independent of one another.

The DRAGEN variant caller applies joint detection (JD) of variants at multiple loci using the following criteria: loci have alleles which overlap each other, loci are in a STR region or less than 10 bases away from an STR region, or loci are less than 10 bases away from each other. STR regions are good candidates for joint detection because 1) this is where PCR-induced indel errors occur, which may overlap with true variant SNV, 2) this is also where true indel variants occur, which may overlap among each other or with SNV, 3) there are situations where a homozygous indel has half of its reads misaligned to represent the indel at the end of a homopolymer rather than at its true location (e.g., beginning or middle of a homopolymer). JD is effective at recovering the true variants in all these cases.

Within JD regions, a haplotype list is generated where all possible combinations of the alleles are represented. Computational complexity increases rapidly beyond a small number of combined loci and events, since it can lead to a large number of haplotypes. In a JD region, the genotyping steps are as follows.

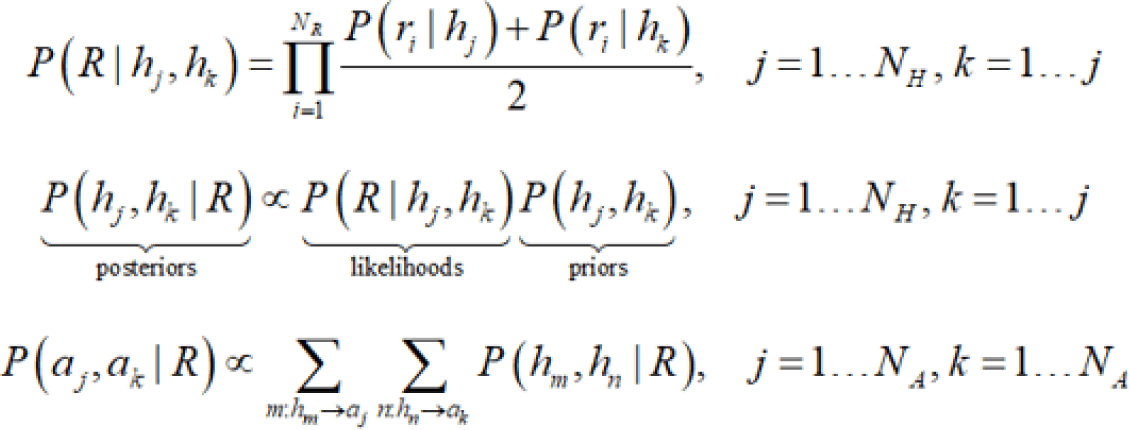

In this case, the posterior probability of diploid combinations of haplotypes over the JD region is computed first, and the pair of events are derived from the most probable pair of haplotypes.

#### Column-wise Event Detection

Legacy haplotype-based variant callers use *de Bruijn* Graph to assemble reads in order to determine candidate haplotypes and identify potential variant sites. But in regions of the genome with tandem repeats, structural variants, and clusters of sequencing errors, the local assembly can fail completely or only give a partial list of candidate haplotypes and variant sites. Local assembly failure can result in lower variant calling sensitivity since we do not genotype the potential variant sites missed by the graph. The DRAGEN variant caller implements a column-wise event harvest scheme that supplements the *de Bruijn* graph by scanning each column of an active region for potential variant sites (SNP and indel) and completes the list of candidate haplotypes with any event found. This restores sensitivity in regions where the graph fails.

#### DRAGEN ML for Germline Small Variant Calling

DRAGEN-ML is a computationally efficient variant calling method with significant gains in False Positive detection. This complements the graph technology approach which primarily helps in recovering False Negatives. Our approach leverages information gathered from the Bayesian variant caller within DRAGEN and refines the called variants in a computationally efficient manner. We also find that ML improves variant calling sensitivity in difficult regions and delivers well calibrated variant call quality scores.

The DRAGEN-ML model is trained using supervised learning, using benchmarks from the GIAB PrecisionFDA dataset^72^(https://precision.fda.gov/challenges/10). We use training data from a range of sequencing platforms and configurations so that the model generalizes well across different sequencer versions, sequencing specific errors (SSEs), lab-preparation flows, coverages, etc.

The machine learning features include statistical descriptions of mapping quality, base quality, strand bias, variant length, GC bias, orientation bias, depth, allele fractions, context, internal HMM scores including foreign read probabilities, SSE triggers, PCR effects, base quality, read position and other statistics from VC internal processing. These features are extracted during DRAGEN variant calling at low computational cost.

The features are used to build a model using offline supervised training, outside the DRAGEN pipeline. The model uses a gradient-boosted ensemble of weak learners to refine variant calls. Given training data, each weak learner partitions the input space using a nonlinear decision tree. Subsequent weak learners are built in a stage-wise fashion to refine the model. The model is trained using gradient descent with regularization and early stopping to address overfitting.

Early stopping holds out a part of the available data as a validation data set. As the model is trained, performance on the validation data is evaluated. Training is terminated when performance on the validation data stops improving – this is an indication that the model is beginning to overfit the training data, impacting generalization capability.

Since DRAGEN-VC has a very high accuracy, only a small minority of calls are false positives, or have genotype errors. This means training data is highly imbalanced. The model is trained using a multiclass cross-entropy loss function that properly models the class probability distribution.

Hyper-parameters (options within model training that affect overfitting and performance, for example learning rate and regularization weights) are tuned using Bayesian optimization. This iteratively refines the hyper-parameters over several training cycles in an efficient manner.

The model classifies each variant call as true, false, or zygosity error, giving corrected genotypes and scores for each variant. The *P(false)* output confidence is used to recalibrate the quality score field. The classifier outputs are used to update the genotype and the genotype quality VCF fields. Potential variants that were scored below threshold by the DRAGEN-VC are recalibrated by ML, and in some cases, this leads to conversion of FNs to TPs. This genotype correction approach leads to reduced numbers of zygosity errors, which are typically counted as both FN and FP, and substantially improves overall performance metrics. The machine learning approach is well calibrated, leading to variant scores that closely match empirical accuracy. The model works well for SNV and for indel.

Subsequently, features generated by variants called within new samples are processed by the pre-trained ML model within DRAGEN-VC, simultaneously with variant calling (**Supplementary Figure S4**). This leads to efficient computation with a low-latency time to answer (additional computation time for DRAGEN-ML is on the order of a minute for a whole genome sample which compares well with the processing time and cost of competing techniques such as deep learning). The predictions from the model are used to update VCF fields, including PL, GP, QUAL, GT and GQ. The QUAL field in the VCF represents the probability of any variant, in phred-scale. GT represents the highest confidence genotype for each variant position. GQ represents the probability of the exact genotype called. PL is a phred scaled likelihood per genotype, and GP is a phred scaled genotype posterior probability.

The machine learning model used in DRAGEN offers global and local interpretation methods, an important feature for users who want to understand why variant calls were made. Other machine learning methods are ‘black-box’ and are harder to interpret.

### DRAGEN Germline Structural Variant Caller

The DRAGEN Structural Variant (SV) Caller is designed to detect structural variants (SV) and indel calls (50bp or larger by default) from Illumina data, such as deletions, duplications, insertions, inversions, and translocations. The process for calling SV in DRAGEN consists of two primary stages. The first stage entails scanning the sequenced genome to collect auxiliary statistics and identify candidate SV regions. These regions are typically either single-locus (for small events) or a pair of loci (corresponding either side of a putative breakpoint). The second stage involves processing the candidate SV regions to identify, refine, score, filter and report SVs. These two stages are similar in principle with Illumina Manta^25^ but DRAGEN SV also includes many unique features and algorithm improvements which result in improved accuracy, precision and run time speed, namely: 1) internal tandem duplication hotspot handling 2) mobile element insertion detection for large insertions; 3) optimize proper pair parameters for large deletion calling; 4) improved assembled contig alignment for large insertion discovery; 5) tuned default parameters/thresholds; 6) refinements in the assembly step; 7) refinements in read likelihood calculations step; 8) improved handling of overlapping mates; 9) improved handling of clipped bases; 10) improved handling of breakpoint homologies; 11) filtering and precision improvements.

One additional feature available in DRAGEN SV v4.2 is the SV multigenome (graph) Hash Table (HT), which can be used as optional on the DRAGEN command line (only the hg38 reference genome is supported in v4.2). The SV multigenome (graph) HT is an augmented version of the default DRAGEN multigenome (graph) HT, which includes common population haplotypes that contain alternative SV alleles. Those SV haplotypes are added to the reference contigs set implicitly to improve alignment accuracy. Read alignments that align best to an SV alternative haplotype carry a “graph alignment” tag (‘ga’ tag which shares the same format as SA tag, but it contains the alignment of the read with respect to the alternative haplotype sequence instead) and are lifted-over (with split mapping if necessary) to the reference contigs accordingly. The DRAGEN SV caller parses the new information provided by the graph reference genome representation in various processing stages. Fully contained reads without a reference liftover are treated as providing support for the alternative SV haplotype. This enables DRAGEN SV to generate full-length assemblies even for insertions larger than twice the library fragment size.

#### DRAGEN SV workflow

In the first stage, a SV candidate graph is created consisting of nodes representing regions of the genome with SV read support between one or more of these loci. SV read support is determined by anomalous read pairs, indels in read alignment, soft-clipped reads, split reads, or ’ga’ tagged reads, which meet specific requirements such as high MAPQ scores. Edges in the graph indicate putative rearrangements between loci, but they do not correspond to specific structural variant hypotheses (such as deletion, insertion, duplication, inversion, or translocation). After a merging support from reads mutually supporting a common SV candidate and denoising to remove poorly supported candidates, the SV candidate region sets are separated into independent structural variant discovery problems and analyzed individually.

Each SV candidate region set identified in the previous step undergoes refinement through local assembly. This involves gathering reads that map to the candidate regions, including from remote regions or a subset of unmapped reads with a ’ga’ tag, expanded by flanking regions.

Reads that support the reference allele, have MAPQ0, are supplementary alignments, or are clipped at both ends by more than 10 bp are filtered out, leaving only the selected reads for assembly. A *de Bruijn* graph approach is used to assemble the selected reads, producing contigs by traversing paths through the graph. The contigs are then scored based on the number of supporting reads, and the highest scoring ones are chosen for further analysis.

High scoring contigs are aligned to reference sequence corresponding to breakend regions using a variant of the Smith-Waterman alignment with a standard affine gap scheme where an additional ‘jump’ state is included to provide a transition between breakend alignment regions if they are distant. The post-assembly alignment process uses features like breakend orientation, breakend distance, alignment quality on flanking regions, and a local breakend depth filter to characterize the SV candidate with the proper SV type and the correct length with single base-pair resolution.

Depending on the type of evidential reads that are associated with each SV candidate, the scoring function in DRAGEN SV will assign weights from the paired-end read component and/or the split read component to the diploid likelihood model ^25^. Filters are then applied in a final step to improve the precision of the scored output.

### DRAGEN Germline Copy Number Variant Caller

The DRAGEN Germline Copy Number Variant (CNV) caller is a read depth and junction based workflow for detection of copy number deletions and duplications. The workflow is defined as a set of subsequent stages, starting from a sample’s input alignments, and ending with an output Variant Call Format (VCF) file containing the inferred Copy Number (CN) alterations for the sample under analysis.

In the first stage, the input reference genome is split into disjoint intervals, with approximately the same size, and the read counts from input alignments falling on each interval are summarized (also denoted as “target [interval] counts”). This approach significantly improves the computation and memory performance of the entire workflow since coverage along the genome is not stored for every base position. The number of bases in each interval depends on its number of *k-mer* unique positions. *k-mer* unique positions are defined as bases where the *k-* mer starting at such position does not show up anywhere else in the genome (default k-mer length is 35 bases). Each interval should have at least n (default: 1000) k-mer unique positions, however, the physical size of the interval cannot grow larger than 2*n. In such cases, the entire interval is discarded, and the interval generation starts from the next genomic position. The use of k-mer unique positions improves precision since it reduces the impact on coverage observed in low-complexity regions due to lower mappability. Finally, for each interval, the “target counts” are summarized as the number of reads falling in such interval that are: primary alignments, not duplicates, properly paired, in forward orientation, with MAPQ ≥ 3, and starting on a k-mer unique position.

Target counts from the previous stage are corrected based on the estimation of GC content in each interval. The purpose of this correction is to reduce GC content related coverage bias. The most likely cause of this bias are PCR artifacts, although GC-bias can also be observed in PCR-free assays. The GC-bias effect on fragment abundance is unimodal: both GC-rich and GC-low intervals are under-represented. GC-bias correction consists of two steps: i) In the measurement step, target intervals are aggregated into different GC content bins. The median read count value for each bin is computed as its correction factor. ii) In the correction step, each correction factor is applied to every read count value within its bin. The target intervals with corrected values are then used in subsequent stages.

The corrected values from each target interval are normalized with respect to the expected baseline level and they will represent copy-ratio values against baseline. The normalization algorithm for Germline CNV is based on the autosomal diploid level from the sample under analysis. Sex chromosomes and pseudo autosomal regions (PAR) regions are handled accordingly based on the sample sex, and any previously excluded intervals are not used for normalization. Briefly, for each contig, the median and total sum of counts are computed. The median divided by the total sum value across all autosomal contigs is then used to compute a distribution of medians. The final median (of the medians’ distribution) is the normalization factor (baseline level) used to compute the copy-ratio values for the sample. The resulting copy-ratio values for each target interval are then transformed in log2-space and median centered.

The next stage is the segmentation stage. The purpose of this stage is to group the input (normalized) target intervals into distinct segments, each segment being assigned a specific state and a normalized log2-transformed copy-ratio. The preferred segmentation mode for the DRAGEN Germline CNV workflow is a variant of the Shifting Levels Model (SLM) ^73,74^. This mode is based on a fixed-state Hidden Markov Model (HMM), which identifies the most likely state of input intervals through the Viterbi algorithm. All states in the HMM emit values following a Gaussian distribution and are specified with equal variance. All states have the same prior probability for the first data point, and state-to-state transition probabilities are the same for all data points. The segmentation output is then post-processed in two steps: i) a segment might be split into sub-segments when some of its targets have different reference ploidy or are separated by a large gap, ii) a segment can be merged with a consecutive one when it contains a small number of bins, and it is not too far from the consecutive one or separated by PAR boundaries. Finally, each segment is summarized with a segment median value (SM), which is computed as the median of its log2-transformed normalized target interval values.

The final stage is the scoring and genotyping stage. The purpose of this stage is to identify the most likely copy number state for each segment and the quality score for the corresponding copy number call. Each copy number state’s coverage is modeled using a Student’s t-distribution with 𝜈 = 30 degrees of freedom. For each segment, the assigned copy number is the one having maximum probability given the segment median (SM), over all possible copy numbers. The maximum likelihood estimate (MLE) of the copy number is emitted in the CN subfield of the genotype field for each segment’s VCF entry. The quality score for each segment call is based on the probability of the directional call (i.e., LOSS, NEUTRAL, or GAIN), rather than specific integer copy number call. This assumes that it is more important to detect the presence of copy number change relative to a reference, than it is to precisely calculate the segment copy number. The Phred-scaled quality score is rounded to the nearest integer and capped at a maximum value. The result is emitted in the QUAL column for each segment entry in the VCF.

When executed in conjunction with the DRAGEN SV caller, the DRAGEN CNV caller extends detection down to 1kbp events by leveraging junction signals. The breakpoint accuracy is improved by split reads and improper pair signals. Depth based calls are reciprocally matched with junction based calls. Previously filtered calls can be rescued if supported by both depth and junction signals and annotated in the VCF file with the SVCLAIM field. This method improves both recall and precision across all length scales.

### ALT masking and hg38 reference improvements

The hg38 reference genome has undergone multiple revisions and improvements over time (**Supplementary Table S17**), as described in this study^24^ and here (https://www.illumina.com/content/illumina-marketing/amr/en_US/science/genomics-research/articles/dragen-demystifying-reference-genomes.html). In the latest DRAGEN v.4.2, we recommend using “hg38-alt-masked-v3” and “hg38-alt-masked-v3-graph” depending on whether DRAGEN multigenome (graph) reference is enabled or not. The improvements between hg38-alt-masked-v3 and the previous hg38-alt-masked-v2 version are as follows:

We included 34 sequences from chm13 and hs37d5 as decoys. Specifically, we included 24 contigs from T2T-CMH13 identified in this study^24^ as well as 4 contigs from hs37d5 identified in these studies ^24,41^ as missing segmental duplication in GRCh38. Furthermore, we included 5 contigs in acrocentric arms of chromosomes 13, 14, 15, and 22 of T2T-CHM13 missing in the GRCh38 assembly and 1 missing duplication of a non-coding region of chr4 identified in chr20 of T2T-CHM13. The main effect of the aforementioned decoys is the variant calling accuracy improvement in the Challenging Medically Relevant Genes FANCD2, MAP2K3, KCNJ18, and KMT2C, as well as in the Y chromosome.

The improvements between hg38-alt-masked-graph-v3 and the previous hg38-alt-masked-graph-v2 version includes the extension of the population samples from 16 samples from European ancestry to 32 samples from different ancestries around the globe. Population SNVs and alt contigs are extracted from the difficult-to-map regions of the genome, extended to include gaps of length <3kbp. Furthermore, we also included one population alt contig from the chromosome 2 of T2T-CHM13 including the TPO gene.

### DRAGEN Gene Specific Callers in Paralog Regions

Segmental duplications account for approximately 5% of the genome. Genes in these regions have traditionally been difficult to interrogate due to high sequence homology. Over multiples releases (**Supplementary Table S6**), DRAGEN has demonstrated that it is possible to variant call some paralog regions by using targeted callers that use fixed base differences gleaned from population level sequencing data to uniquely and correctly place reads to call CNV, SNV, and indels. These callers can be run as part of the regular DRAGEN workflow with no meaningful run time increase and share similar subcomponents.

First, the total copy number of the gene and its paralog is computed from the counts of reads aligned to regions in either the gene or its paralog. A series of pre-selected differentiating sites across the gene and paralog regions are then used to identify the gene-specific copy number for various segments of the gene. These differentiating sites were selected at positions with sequence differences between the gene and paralog. Structural variant calling is performed from the gene-specific copy numbers across the gene to detect various hybrid structures between the gene and paralog, optionally including phasing of the various detected haplotypes to detect gene conversion events. Following structural variant calling, small variant calling may also be performed at a set of pre-determined sites containing known variants in the population. For variants occurring in homologous regions of the gene, a joint analysis of reads mapping to either the gene or the paralog is performed since reads containing these variants may map to either location in the reference genome.

The methods used for each targeted gene caller are given below.

#### CYP2D6 caller

The CYP2D6 caller (https://www.illumina.com/content/illumina-marketing/amr/en_US/science/genomics-research/articles/PGx-research-blog.html) identifies the total copy number of the gene-pseudogene pair, as well as for a 1.6 kbp tract of unique sequence adjacent to *CYP2D7*. This unique region co-occurs with *CYP2D7* and *CYP2D6-CYP2D7* fusion genes, so the total copy number of *CYP2D6* and *CYP2D6-CYP2D7* fusion genes can be found by subtracting the copy number of the unique region from the copy number of the gene-pseudogene pair. Fusion genes are further identified by scanning across differentiating sites within the gene-pseudogene pair, calculating the gene copy number at each differentiating site, and finding changes in copy number states that indicate where a gene fusion has occurred.

Small variants are identified by read analysis. In regions of the gene where reads can be confidently aligned to CYP2D6, reads from only that gene are considered. In regions where high homology makes confident alignment impossible, reads from either gene are used to call variants without phasing to gene or pseudogene.

#### SMN1/2 caller

The *SMN1/2* ^27^ caller identifies the total copy number of the *SMN1-SMN2* pair using reads aligned to a 22kbp region, which includes exons 1-6. The copy number of intact (non-truncated) *SMN* is determined by calling the copy number of a 6kbp region (with exons 7-8) and subtracting that copy number from the total. If there is a difference, a truncation is identified.

A scan is then performed across the differentiating sites to determine the most likely *SMN1* vs *SMN2* copy number at each site, and the resulting site copy numbers are combined to generate a consensus *SMN1* copy number. *SMN1* copy number is found by subtracting *SMN1* copy number from total copy number.

#### GBA caller

The *GBA*^28^ caller identifies the total *GBA-GBAP1* copy number, as well as the copy number of a 10kbp unique region between them. Non-diploid copy number of this region indicates that a recombinant variant has occurred. Copy number of less than two indicates a deletion, while more than two indicates a duplication. The breakpoint of recombinant variants is then identified. First, a scan is performed across differentiating sites and the copy number of the GBA allele is calculated at each. Transitions from one copy state to another between pairs of differentiating sites indicate the breakpoint region. The breakpoint is further refined by performing read-based phasing across the 1.1kbp high-homology region, using differentiating sites between tracts of identical sequence as evidence. Haplotypes are identified which convert from *GBA* to *GBAP1* alleles, indicating the breakpoint location.

Small variants are identified by read analysis. In regions of the gene where reads can be confidently aligned to the gene, reads from only the gene are considered. In regions where high homology makes confident alignment impossible, reads from either gene are used to call variants without phasing to gene or pseudogene.

#### CYP2B6 caller

The *CYP2B6* caller (https://www.illumina.com/content/illumina-marketing/amr/en_US/science/genomics-research/articles/PGx-research-blog.html) identifies the total copy number of *CYP2B6* and *CYP2B7*. Small variants are identified by read analysis. In regions of the gene where reads can be confidently aligned, reads from only the target gene are considered and variant calls assigned uniquely to the gene location. The homology-region gene conversion variant is called by a dedicated method. Reads that align to that location, whether from the gene or from the pseudogene, are tested for the pathogenic allele. If found, a small number of nearby differentiating sites are employed in read-based phasing and site-by-site read depth analysis. These pieces of evidence indicate if the pseudogene allele occurs on the same haplotype as gene alleles at nearby differentiating sites, indicating a gene fusion/gene conversion.

Structural and small variants identified are matched against 39 known star allele definitions. The two haplotypes with highest likelihood (based on variant copy number and population frequency in the 1kGP) are selected for reporting. If no variants called match known star alleles, no call is reported. If multiple genotypes are identified with similar population frequencies, all genotypes are reported.

The accuracy of the CYP2B6 caller was assessed against 125 samples from 1KGP and 76 Coriell samples. Calls on the Coriell samples were compared against either GeT-RM or calls from Stargazer. For the six samples where calls were not concordant with GeT-RM, DRAGEN was concordant with Stargazer on five samples and Stargazer did not make a call on the sixth sample. Calls on the 125 samples from 1kGP were compared against calls manually curated from long-read sequencing data. **Supplementary Table S6** shows the concordance results of the DRAGEN and CYP2B6 caller.

#### HLA caller

The HLA caller in DRAGEN is specifically designed for genotyping HLA-A and HLA-B genes, which are highly polymorphic; that is, these genes have thousands of defined haplotypes across the population, each containing hundreds of variants. This caller uses expectation maximization to analyze reads aligning to full sequence alleles from the IMGT/HLA database and Allele Frequency Net Database to output two-field resolution of HLA.

The DRAGEN v4.0 HLA caller was validated for genotyping accuracy with 117 WGS samples from the 1000 Genomes consortium. Of the 351 calls (three genes for each of the samples), 349 calls were concordant with Sanger sequencing results (accuracy 99.6%).

#### HBA1/2 caller

The *HBA1/2* caller (https://www.illumina.com/content/illumina-marketing/amr/en_US/science/genomics-research/articles/HBA-targeted-caller.html) identifies the total copy number of *HBA1* and *HBA2*, and the copy number of a region around them. This is used to determine, in the case of a copy number variant, the copy number genotype of *HBA1* and *HBA2* genes. Pathogenic small variants are identified using reads aligned to either gene and variants are reported without phasing to gene or pseudogene.

#### CYP21A2 caller

Total copy number of *CYP21A2* and *CYP21A1P* is determined using a region of the segmental duplications that includes the entire target gene but omits other common copy number variants nearby. Small variants in high-homology regions are identified using reads from both gene copies, while *CYP21A2* reads are used to test for a small number of variants in unique regions.

The frequency of recombination between segmental duplication copies calls for targeted identification of gene conversion variants, as in *GBA*. A set of differentiating sites across most of the *CYP21A2/CYP21A1P* genes are used in read-based phasing to identify case switches, from pseudogene to gene, in the same haplotype. These are reported as recombinant variants.

#### RHD/RHCE caller

The RHD/RHCE caller calls the RHCE*CE-D(2)-CE gene conversion. It identifies the total combined RHCE/RHD copy number using reads aligning to either gene. It then scans across a set of differentiating sites within the genes to determine the copy number state of each gene at each site. Identification of a copy state change in consecutive differentiating sites shows evidence of the gene conversion event. Both RHD->RHCE and RHCE->RHD must be detected for a call. Read-based phasing is then performed across the region of the gene conversion to support the call and refine the breakpoint location, using the differentiating sites as evidence.

The breakpoint can be refined as phasing reveals haplotypes with transition from RHD-only alleles to RHCE-only alleles or the reverse. In cases where haplotypes indicate gene conversion from one gene to the other and then back, the differentiating sites that provide evidence for the gene conversion are reported as variant sites.

#### LPA caller

The LPA^29^ caller identifies the total KIV-2 copy number using reads aligned to any of the six copies of the repeat unit in the reference genome. That reference copy number is used as a special scaling factor to determine the number of copies of KIV-2 in the sample, rather than copies of the six-copy reference sequence.

The effect of KIV-2 on transcript length means that phased/per-haplotype copy number is also an important factor to determine. Two marker sites within the repeat unit can be leveraged for this. These sites, identified by diverse trio analysis within the 1kGP, are polymorphic but consistently have the same allele within each same-haplotype copy of KIV-2. Reads are therefore collected at these sites, from any copy of the KIV-2 repeat, and the reference/alternate alleles counted. If both reference and alternate marker site alleles occur, the sample is considered heterozygous for the markers. The reference-allele read fraction is then used as a multiplier for total copy number to determine the number of KIV-2 copies with the reference marker allele. The same analysis using alternate marker allele fraction determines the number of copies with the alternate allele, and phased allelic copy number is found. As this analysis requires the heterozygous state for the marker sites, it is possible in a subset of genomes, averaging ∼50% across the 1kGP.

### Iterative gVCF Genotyping beyond million sample size scale

In multi sample VCF (msVCF), all the variants called at cohort level are stored, with all samples genotyped at every variant site. The overarching challenges in this joint analysis include variant quality, performance and scalability, and solving the N+1 problem (i.e., iteratively aggregating a new batch of samples with the existing batches, without reanalyzing the existing ones).

The DRAGEN Iterative gVCF Genotyper (IGG) can efficiently aggregate hundreds of thousands to millions of gVCFs from the DRAGEN germline variant calling pipeline, perform joint calling and genotyping, and generate a msVCF file. The output msVCF file also contains cohort level variant statistics (including allele frequency, sample genotype rate, coverage rate) and QC metrics (such as Hardy Weinberg test p-value, and inbreeding coefficient) that can be used for downstream variant filtering.

IGG splits the samples into batches (e.g., 1000 samples per batch) and genome into shards (1% of genome per shard). The process consists of 3 steps, and each step is highly parallelized by batch and by shard across distributed compute nodes (minimum 16 cores, 32G memory).

For variant comparison we utilize the start location or precise reported position per variant to compare them.

In step1, for each batch, gVCFs files are aggregated into a customized data format, aka Cohort files, which store the compressed sample level metrics data. The variant statistics are stored in another customized data format, aka Census files. In step 2, Census files from all batches are aggregated into a Global Census file, which stores the global variant statistics and normalized variant alleles. In step 3, for each batch, msVCFs are generated for all the variants called in the Global Census file and sample level metrics are retrieved from the batch Cohort and Census files.

In N+1 scenario, IGG minimizes the cost of recompute, by requiring only new batches to be aggregated in step 1, followed by a quick update of Global Census from both old and new batches in step 2, and output of msVCFs for both old and new batches in step 3. This is achieved since in the Cohort files, we use hash compression to store per sample variant metrics in hash tables and then encode them into a generic htslib^75^ bed file format, so that the tabix indexing allows random access based on position. We cluster gVCFs records in all batch samples by regions for both variants or hom-ref records.

IGG addresses the limitation of storing millions of samples by compressing the sample metrics into a localized format (e.g., LPL, LAD). The batch/shard data partition scheme also allows for highly parallelized downstream variant analysis. The msVCFs from different batches contain the same number of variants, allele order and global variant statistics, making it straightforward to merge across multiple batches, and concatenate into chromosomes.

### Illumina Connected Annotations (ICA)

In preparation for usage, Illumina Connected Annotations (ICA) performs re-structuring and compression of data from annotation sources (**Supplementary Table S17**) into pre-computed caches for highly efficient parallelized querying, analogous to preparing a reference genome for DRAGEN mapping and alignment. This can be repeated to provide updated annotation content independent of the annotations software version. The annotations software reads in single or multi-sample VCFs for small variants (including MNVs), CNVs, SVs, and/or STRs, such as those generated by the DRAGEN DNA analysis pipeline.

In the first phase of analysis, all variant types are annotated using reciprocal overlap with cytobands and known CNVs/SVs from interval-based annotation sources. Next, the software computes sample-specific variant allele frequencies, ensuring availability of these critical values not always present in the input VCFs.

In the second phase, the annotations software refines each alternate allele to its most parsimonious representation. The HGVS^76^ genomic notation applies a right-aligned approach to this representation, while all other annotations follow a left-aligned format per NGS conventions. Following this refinement, alternate alleles are matched to variant databases such as gnomAD and ClinVar. Exact allele matching is required for population frequency data. For other sources such as ClinVar, both exact and overlapping matches are included and marked accordingly. ICA then applies repeat size thresholds to classify STR variants as “normal” or “expanded” based on user-defined thresholds or a default threshold set as described previously^23^

In the third phase, 1) identifies transcripts intersecting each alternate allele using an interval array, 2) marks overlapping exons and/or introns, 3) adjusts for discrepancies between transcript and genomic reference sequences, and 4) provides predicted impact on coding sequence (“c.” or “cNomen”) and protein sequence (“p.” or “pNomen”). Canonical transcripts are identified using information from MANE^77^ or, when not available, via existing heuristic methods^78,79^. This phase also provides consequences relevant to each variant using Sequence Ontology^80^ standard nomenclature. ICA performs right-alignment to coding and protein sequences as needed, in accordance with HGVS standards^76^. If applicable, it also adds associated cancer hotspot annotation. Then it evaluates SVs to identify potential unidirectional gene fusions based on the resulting gene orientation and proximity. Known fusions are annotated using paired gene symbols from resources such as COSMIC and FusionCatcher. In the final phase, it adds gene-specific annotations for each unique gene with at least one variant in the VCF, retrieving data from OMIM, ClinGen, and other gene information sources.

The output of ICA is a structured, indexed, and compressed file in JSON format that can be queried directly using JASIX (an included tool analogous to TABIX for VCF manipulation) or used as input for downstream tertiary analysis platforms such as Illumina Connected Insights. ICA utilizes an interval array data structure to optimize for speed and a comprehensive testing system to ensure accuracy, thus suiting the demanding requirements of population-scale WGS.

The structure and function of ICA offer several key advantages. First, the highly compressed binary data files, interval arrays, and multi-sample inputs, enables it to annotate a single human genome (roughly 4-6 million variants) within 12 minutes using a DRAGEN server. Second, finely tuned alternate allele refinement (the normalization and left-alignment of each alternate allele mentioned in the second phase) and transcript corrections (adjusting for the discrepancies between the transcript and genomic reference sequences mentioned in the third phase) greatly reduce the number of erroneous consequence predictions and missed annotations. To account for the nuances of both the HGVS ruleset and imperfect transcript-to-genome mapping, ICA utilized the BioCommons hgvs package^81^ to create an extensive test suite containing more than 18 million variants to achieve 99.995% accuracy in HGVS c. notation and 99.986% accuracy in HVGS p. notation. Finally, the breadth of supported inputs and variant types streamlines WGS workflows that would otherwise require different annotation tools for each variant type.

### Variant calling comparison and benchmarking

Mapping and Variant calling:

For the DRAGEN end-to-end variant calling pipeline, the illumina NovaSeq 6000 PCR-free 35x sequencing of all samples were uploaded to illumina’s ICA platform where the alignment and variant calling was performed using the DRAGEN 4.2 pipeline. The command and parameters used for the DRAGEN run are given below.

`dragen \`

`--ref-dir <path-to-hg38-alt_masked.graph.cnv.hla.rna_v3> \`

`--fastq-file1 <PATH-to-R1-fastq> \`

`--fastq-file2 <PATH-to-R2-fastq> \`

`--enable-map-align true \`

`--enable-map-align-output true \`

`--output-format CRAM \`

`--enable-duplicate-marking true \`

`--enable-variant-caller true \`

`--vc-emit-ref-confidence GVCF \`

`--vc-enable-vcf-output true \`

`--enable-cnv true \`

`--enable-sv true \`

`--vc-ml-enable-recalibration true \`

`--repeat-genotype-enable true \`

`--repeat-genotype-use-catalog expanded \`

`--enable-targeted true \`

`--enable-pgx true \`

`--cnv-enable-self-normalization true \`

`--intermediate-results-dir /scratch \`

`--output-file-prefix <SAMPLE-name> \`

`--output-directory <OUTPUT-path-directory> \`

`--force`

The above command performed SNV and indel calling including ML recalibration, CNV calling, SV calling, STR calling, and targeted calling.

For the BWA based variant calling pipelines, first the illumina NovaSeq 6000 PCR-free 35x sequencing of all samples are mapped using BWA (v0.7.15) (with parameters -K 100000000 -Y -t 8 -R @RG\tID:0\tSM:HG002\tLB:HG002\tPU:HG002_38_nodecoy\tCN:BCM\tDT:2021-03-10T00:00:00-0600\tPL:Illumina) to both GRCh37 and GRCh38 reference genome. The GRCh37 reference is used because the SV benchmark set is only available for that reference. Following is one of the commands used for mapping HG002 dataset to the GRCh38 reference. For SNV and indel calling, we used GATK (v4.2.5.0) Haplotypecaller with `--java-options “-Djava.io.tmpdir=${TMP} -Xms20G -Xmx20G` parameters. We also run DeepVariant (v1.5.0) using singularity pull `docker://google/deepvariant:”${BIN_VERSION}”` and performed the singularity run with the GRCh38 reference and alignment i.e., BAM files generated using BWA-MEM v0.7.15. The following singularity was used for the HG002 dataset.

`singularity run \`

`--bind “${INPUT_DIR}:/mnt/input,${REF_DIR}:/mnt/reference,${OUTPUT_DIR}:/mnt/outp ut,${BIND_TMPDIR}:/tmp” \`

`deepvariant_1.5.0.sif \`

`/opt/deepvariant/bin/run_deepvariant \`

`--ref=“/mnt/reference/hg38.fa” \`

`--reads=“/mnt/input/${SAMPLE}_hg38_sorted.bam” \`

`--model_type=“WGS” \`

`--sample_name=“${SAMPLE}” \`

`--output_vcf=“/mnt/output/${SAMPLE}.vcf.gz” \`

`--output_gvcf=“/mnt/output/${SAMPLE}.g.vcf.gz” \`

`--num_shards=“1”`

For SV calling, we used Manta (v1.6), Delly (v1.16), and Lumpy (v0.3.1) with their default parameters given the bam file from BWA-MEM v0.7.15 (GRCh37 reference). The SV calling by Lumpy first needs pre-processing to extract the discordant read-pairs (using `samtools view-b -F 1294)` and the split-read alignments using samtools and the customized script extractSplitReads_BwaMem that is provided with the tool. After these steps, we run the `lumpyexpress` executable with the original BAM file, the split-read alignment BAM and the discordant read-pair BAM as inputs and all other default parameters. For Delly, we converted the generated BCF file to VCF file using bcftools (v1.15.1).

For CNV calling, we used CNVnator (v0.4.1) with default parameters in addition to the DRAGEN 4.2 pipeline on ICA.

For the Giraffe based pipeline, we followed the WDL pipeline as specified in (https://zenodo.org/record/6655968#.ZHYsCy_MKgQ), using the minaf.0.1 GRCh38 reference released on AWS (https://s3-us-west-2.amazonaws.com/human-pangenomics/index.html?prefix=pangenomes/freeze/freeze1/minigraph-cactus/filtered/). We aligned the reads using Giraffe v1.48.0. The command lines and parameters are as follows.

`vg giraffe \--progress \`

`--read-group “ID:1 LB:lib1 SM:HG002 PL:illumina PU:unit1” \`

`--sample “HG002” \`

`--prune-low-cplx \`

`--max-fragment-length 3000 \`

`--output-format bam \`

`-f <PATH-to-R1-fastq> \`

`-f <PATH-to-R2-fastq> \`

`-x hprc-v1.0-mc-grch38-minaf.0.1.xg \`

`-H hprc-v1.0-mc-grch38-minaf.0.1.gbwt \`

`-g hprc-v1.0-mc-grch38-minaf.0.1.gg \`

`-d hprc-v1.0-mc-grch38-minaf.0.1.dist \`

`-m hprc-v1.0-mc-grch38-minaf.0.1.min \`

`-t 32 > HG002.giraffe.grch38.minaf.0.1.bamsort the output BAM with sambamba v0.8.1 and index with samtools v1.15.1`

`sambamba sort \-t 32 \`

`-o HG002.giraffe.grch38.minaf.0.1.sort.bam \ HG002.giraffe.grch38.minaf.0.1.bamsamtools index \`

-@ 32 \ HG002.giraffe.grch38.minaf.0.1.sort.bamleft shift using FreeBayes v1.20

`bamleftalign < HG002.giraffe.grch38.minaf.0.1.sort.bam \> HG002.giraffe.grch38.minaf.0.1.sort.left.shifted.bam \`

`--fasta-reference hprc-v1.0-mc-grch38-minaf.0.1.fa \`

`--compressedIdentified targets for indel realignment using GATK v3.8.1 and bedtools v2.21.0`

`java -jar GenomeAnalysisTK.jar -T RealignerTargetCreator \--remove_program_records \`

`-drf DuplicateRead \`

`--disable_bam_indexing \`

`-nt 32 \`

`-R hprc-v1.0-mc-grch38-minaf.0.1.fa \`

`-I HG002.giraffe.grch38.minaf.0.1.sort.left.shifted.bam \`

`--out HG002.giraffe.grch38.minaf.0.1.sort.left.shifted.intervalsawk -F ’[:-]’ ’BEGIN { OFS = “\t” } { if ($3 == “”) { print $1, $2-1, $2 }`

`else { print $1, $2-1, $3}}’ HG002.giraffe.grch38.minaf.0.1.sort.left.shifted.intervals > HG002.giraffe.grch38.minaf.0.1.sort.left.shifted.intervals.bed && \`

`bedtools slop -i HG002.giraffe.grch38.minaf.0.1.sort.left.shifted.intervals.bed \-g hprc-v1.0-mc-grch38-minaf.0.1.fa.fai \`

`-b 160 >HG002.giraffe.grch38.minaf.0.1.sort.left.shifted.intervals.widened.bed`

`Indel realign using Abra v2.23`

`java -Xmx16G -jar abra2-2.23.jar \`

`--targets HG002.giraffe.grch38.minaf.0.1.sort.left.shifted.intervals.widened.bed \--in HG002.giraffe.grch38.minaf.0.1.sort.left.shifted.bam \`

`--out HG002.giraffe.grch38.minaf.0.1.sort.indel.realigned.bam \`

`--ref hprc-v1.0-mc-grch38-minaf.0.1.fa \`

`--index \`

`--log warn \`

`--threads 32Variant calling using DeepVariant v1.5.0 with the following singularity command.`

`singularity run \`

`--bind “${INPUT_DIR}:/mnt/input,${REF_DIR}:/mnt/reference,${OUTPUT_DIR}:/mnt/outp ut,${BIND_TMPDIR}:/tmp” \deepvariant_1.5.0.sif \`

`/opt/deepvariant/bin/run_deepvariant \`

`--ref=“/mnt/reference/hprc-v1.0-mc-grch38-minaf.0.1.fa” \`

`--reads=“/mnt/input/HG002.giraffe.grch38.minaf.0.1.sort.indel.realigned.bam” \`

`--model_type=“WGS” \`

`--sample_name=“HG002” \`

`--output_vcf=“/mnt/output/HG002.vcf.gz” \`

`--output_gvcf=“/mnt/output/HG002.g.vcf.gz” \`

`--make_examples_extra_args=min_mapping_quality=1 \`

`--num_shards=“1”`

#### Filtering and counting

Only the variants with PASS filter and non-REF calls (i.e., the ALT is not “.”) are retained for further analysis. We used the `bcftools stats` command to count SNV and indel variants. For the SV VCF files, the inversion (INV) and translocation (TRA) variant types are marked as SVTYPE=BND, so we used a customized script (https://github.com/srbehera/DRAGEN_Analysis/blob/main/convertInversion.py) that changes the SVTYPE value of inversion types from BND to INV e.g., SVTYPE=INV using the following commands.

`python2.7 convertInversion.py <SAMTOOLS_PATH> <REF_PATH> <VCF_FILE>`

The remaining BND types are considered to be TRA types. The actual number of TRA types is counted by the counting of BNDs and match MATE_BNDs then divide them by 2. The counting of other variants was done by just counting variants with SVTYPE=<VARIANT_TYPE> where Variant_type is either INS or DEL or DUP or INV.

#### Benchmark

The benchmarking of variants was performed using the GIAB benchmark set for both small variants and structural variants.

For small variants we benchmarked each of the SNV VCF files once with the genome wide benchmark (GIAB v4.2.1) and once with the challenging medically relevant genes (GIAB v1.0)^12^. This was performed for HG001-07 to assess the variant performance on all available samples (https://ftp-trace.ncbi.nlm.nih.gov/ReferenceSamples/giab/release/). For evaluation we used the `vcfeval`^82^ option of RTGTools (v3.12.1) with parameters `-m roc-only` along with other inputs e.g., benchmark set `(-b),`high-confidence bed regions for benchmark set `(--bed-regions)`, SNV VCF file`(-c)`, reference sequence formatted to SDF format `(-t)`and reported the values based on PASS filter. For generating the SDF format of reference sequence, we used `format` option of RTGTools.

For the Structural Variant calling benchmark we compared the obtained insertion and deletions and compared it to the GIAB benchmark (v0.6) on GRCh37. In addition to genome wide we also benchmarked the CMRG benchmark for SV (v0.6) ^39,82^. We evaluated all the SV call sets based on HG002 only using Witty.er (v0.3.5.1) with default config file provided in github repo (https://github.com/Illumina/witty.er) and `-em SimpleCounting` parameters. The following command is used for running the Witty.er.

`Wittyer.dll -i <INPUT_VCF> -t HG002_SVs_Tier1_v0.6.vcf.gz`

`--includeBed HG002_SVs_Tier1_v0.6.bed --configFile config_wittyer.json - em SimpleCounting -o <OUT_FILE>`

For CNV calls, we could only evaluate deletions as there are no duplications reported on HG002 benchmarks. We compared the results to deletion calls from GIAB SV benchmark (v0.6) for GRCh37 that are 1Kbp or larger (using SVTYPE=DEL and SVLEN <= -1000 filters). This was again evaluated using Witty.er (v0.3.5.1) with `-em CrossTypeAndSimpleCounting` parameter and all other default parameters.

For STR discovery, DRAGEN was run with `–-repeat-genotype` parameters and a catalog of approximately 50K regions and 174K regions. GangSTR (v2.5) was run with the catalog (https://s3.amazonaws.com/gangstr/hg38/genomewide/hg38_ver13.bed.gz) provided on their Github repository. The following command was used to run GangSTR. The BAM file is generated by aligning the HG002 NovaSeq 6000 PCR-free 35x sequences to the NCBI GRCh38 reference.

`GangSTR --bam HG002_hg38.bam`

`--ref GCA_000001405.15_GRCh38_no_alt_analysis_set.fasta`

`--regions hg38_ver13.bed`

`--out<OUTPUTPREFIX>`

For benchmarking, the VCFs generated by both DRAGEN and GangSTR were converted into VCF4.2 specifications by using custom scripts (see **Data Availability**). The evaluations were performed using the Truvari (v4.1-dev) and the GIAB benchmark VCF and bed regions (https://github.com/ACEnglish/adotto/tree/main). Truvari performs the evaluation in two stages:

1. benchmarking using `truvari bench` 2) refinement using `truvari refine`. Following is the command used for the first stage.

`truvari bench -b GIABTR.HG002.benchmark.vcf.gz \`

`-c <VCF> \`

`--includebed GIABTR.HG002.benchmark.regions.bed.gz \`

`--sizemin 5 --pick ac -o bench_result/`

For the refinement stage, three different approaches were used. First, the refinement was performed using the GIAB bed regions only.

`truvari refine --use-original-vcfs --reference ${ref} bench_result/`

Then, the bed regions used by callers make sure the individual callers are not penalized for the regions that are outside of individual bed regions. The output file `refine.region_summary.json` contained the evaluation results.

`truvari refine --use-original-vcfs --reference ${ref} --regions`

`<individual_regions.bed> --align mafft bench_result/`

Finally, to make a comparison of STR calls in the region that are common to DRAGEN and GangSTR, we used bedtools intersect of two bed regions and then used the refinement commands of truvari.

`truvari refine --use-original-vcfs --reference ${ref} --regions`

`<intersect.bed> --align mafft bench_result/`

#### Mendelian inconsistency

The mendelian plugin of `bcftools (v1.15.1)` was used for detecting the mendelian inconsistency. The multi-sample VCF (msVCF) files for the trios were generated using DRAGEN. The following commands are used.

AshkenazimTrio: `bcftools +mendelian HG002_HG003_HG004.vcf.gz -t PrecisionFDA_v2_HG004_hg38,PrecisionFDA_v2_HG003_hg38,PrecisionFDA_v2_HG00 2_hg38`

ChineseTrio: `bcftools +mendelian HG005_HG006_HG007_GIAB.vcf.gz -t HG007_NA24695_Mother_HiSeq_40x,HG006_NA24694_Father_HiSeq_40x,AWS_HG005_40 x_hg38`

For GIAB high confidence regions, we extracted the variants form the msVCF that intersects with the GIAB (v4.2.1) HG002 high confidence regions and then used similar commands as given above for mendelian inconsistency.

For De Novo variant rate computation was done by counting the variants with “DeNovo” tag in the msVCF files.

#### 1kGP small variant analysis

The individual small variant VCF files of DRAGEN runs were combined to multi-sample VCF file using DRAGEN’s Iterative GVCF Genotyper Analysis platform that works on three steps: a) gVCF aggregation b) Census aggregation and c) msVF generation. The first step aggregates the batch of gVCF files into a Cohort and a Census file. The cohort file stores the gVCF data of multiple samples in a condensed format and the census file stores summary statistics of all the variants and hom-ref blocks among samples in the cohort. The second step creates a census file of all samples taken together. Finally, the last step generates a multisample VCF containing the variants and alleles discovered in all samples from all batches, and also includes global statistics such as allele frequencies, the number of samples with or without genotypes, and the number of samples without coverage.

The multi-sample VCF files are first left-aligned and normalized with bcftools(bcftools norm `-f ${ref} -m -both).` For the DRAGEN callset, the variants with GT = “, QL=., DP = 0 or ., GQ = . are considered as “no data” (in both DRAGEN and GATK msVCFs). The variants with GT=./. Or ALT=NON_REF are considered as “no genotype”. The variants with “no data” or “no genotype” and zero allele count (AC=0) are filtered. For the GATK callset, the variants with ALT=* are also filtered. For both GATK and DRAGEN callset, only variants with filter=PASS is considered for further analysis. The number of variants is computed at cohort level as well as sample level (averaged by population).

For allele frequency (AF) based analysis and finding the known and novel variants, the variants are annotated using ICA, a variant annotation pipeline for genomic variants. We annotated small variants called from a subset of 2504 unrelated samples, by extracting site level VCF from the multi-sample VCF as the input of the annotation pipeline. From the annotation result, we define novel variants as those do not present in dbSNP (build 155). For each variant the functional annotation is retrieved from the transcripts consequence of the ICA output JSON file. The count of variants was generated for both known and novel based on three AF bins (singleton, rare: AF <= 1% and common: AF > 1%).

#### 1kGP large variant analysis

For the analysis of large variants (>= 50bp) generated by DRAGEN for the 1kGP cohort, we first merged the STR, CNV and SV VCF files of each individual independently by first splitting multi-allelic sites into separate VCF entries using the normalization command of bcftools (v1.15.1).

We then collapsed redundant calls between type representations using a custom script (dragen_sv_merge.py) which leverages the Truvari (v4.1.0) api^50^. This script identifies redundant variant representations between STR and SV VCFs as well as redundancies between SV and CNV VCFs before outputting a single, unified VCF. To be considered redundant, SV representations up to 500bp in length with at least 70% size similarity to an overlapping STR representation of matching type is removed. Here, matching type is defined as SV deletions being synonymous to STR contractions and SV insertions to STR expansions.

Similarly, CNV representations with at least 70% size similarity and within a maximum distance of 1kbp of an SV representation at least 800bp in length and of matching type is removed. Here, matching type is defined as SV and CNV deletions or SV insertions and duplications with CNV duplications. In the final merging step, a project-level VCF is produced using `bcftools merge` to consolidate genotypes from identical variants between samples. The resulting project level VCF is then further normalized to ensure variant representations’ reference alleles have a consistent adherence to VCF format specification using `bcftools norm --check-ref s --fasta-ref`. Finally, we filtered variants in centromeric, pericentromeric etc. regions and generated the final SV callset. We counted the STRs and CNVs in the final merged file using the STR and cnv tags and the remaining variants were counted as SVs. The number of different SV types (DEL, INS, DUP, INV, TRA) were counted using SURVIVOR^83^ (v1.0.6). We also generated the SV counts for different types per individual using SURVIVOR and computed the average counts for super-population. The allele frequency of variants was calculated using VCFtools (v0.1.6) for all SVs separately.

For the PCA plot analysis, we first extracted the SVs at chromosome level merged and normalized multi-sample VCF files with MAF >= 0.05 using bcftools (v1.15.1) and then used principal component analysis module of AKT (v0.3.3) (https://github.com/Illumina/akt) on the extracted variants using the following command.

`akt pca file.vcf.gz --force -Oz -o file.pca.cf.gz > file_pca.txt`

For variant annotation, we used the merged variant file that is normalized and the multi-allelic sites split into different lines. We extracted the variants for 2,504 unrelated samples and then the annotation was done using Illumina Connected Annotations (ICA) from three different sources: gnomAD, 1kGP and TOPMed^19,84^. The novel variants are the ones with counts for all these four sources marked as zero in the annotated VCF file and the remaining i.e., with count > 0 for at least one source is considered to be known variant. The Pearson correlation coefficient and p-value calculations for allele count and allele frequency were calculated using `pearsonr` function from the `scipy.stats` module of `numpy(v1.25)` python library.

The overlapping SVs with exon, intron and intergenic regions were extracted using bcftools^75^ and corresponding bed regions extracted from Genecode annotation file (release 43).

## Data availability

The DRAGEN VCF files and trageted caller JSON files were uploaded to https://zenodo.org/uploads/8350256

The SV and CNV VCF files (GRCh37 reference with DRAGEN, Lumpy, Manta, Delly, CNVNator), SNV VCF files (GRCh38 reference with GATK+BWA and DeepVariant+BWA and DeepVariant+Giraffe) were uploaded to https://zenodo.org/uploads/10428664.

AWS bucket for 1kGP DRAGEN4.0 : https://s3://1000genomes-dragen-v4.0.3/data/cohorts/gvcf-genotyper-dragen-4.0.3/hg38/3202-samples-cohort/

minaf.0.1 GRCh38 reference https://s3-us-west-2.amazonaws.com/human-pangenomics/index.html?prefix=pangenomes/freeze/freeze1/minigraph-cactus/filtered/)

## Code availability

Scripts used in this study https://github.com/srbehera/DRAGEN_Analysis . DRAGEN v4.2 is freely available for academic institutions upon request.

## Acknowledgment

We would like to thank AWS for generous compute support for the 1000 genome project data set. FJS, SB, AE are partially supported by NIH grants (1U01HG011758-01)

## Contributions

FJS, SC, and RM designed the project. SB, MR, ST, ZH did the experiments and all the analysis. All the authors reviewed and edited the manuscript.

